# Potassium ferric oxalate nanoparticles prevent human blood clotting and thrombosis in a mouse model

**DOI:** 10.1101/2024.11.04.621820

**Authors:** Devyani Yenurkar, Ankit Choudhary, Anoushka Shrivastava, Pragya, Snehasish Mandal, Priyanshu Soni, Lipi Pradhan, Ankur Singh, Arnab Sarkar, Sudip Mukherjee

## Abstract

Blood clot creates occlusion in the veins and arteries, which leads to pernicious effects. Here, the anticoagulation properties of potassium ferric oxalate nanoparticles (KFeOx-NPs) in human blood were demonstrated for blood clot management. The mechanism involves the chelation of calcium ions from the blood by the oxalate present in the KFeOx-NPs. Various commercial assays were used to determine the clotting time for the KFeOx-NPs and identified the hindrance in activating factor XII in the intrinsic pathway. We used animal models to show toxicity and biodistribution profiles and determined the safety and efficacy. Intravenously injected KFeOx-NPs increased clotting time and thrombosis prevention in a mouse model confirmed by ultrasound and the power Doppler images. Coating catheters with KFeOx-NPs prevents clot formation with reduced protein attachment when incubated with blood, enhancing blood flow properties. In biological applications, KFeOx-NPs may improve the long-term prevention of blood clot formation and enhance the efficiency of medical devices.

**TOC:** 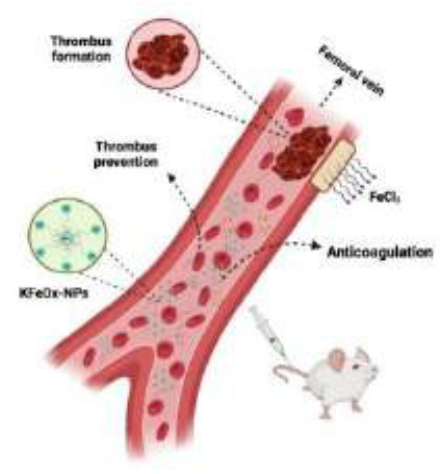

PVP-stabilized novel potassium ferric oxalate nanoparticles were synthesized for the application of blood clot management and thrombosis prevention.

## 1. Introduction

Blood clot-related diseases have one of the highest rates of mortality among non-communicable diseases.^[1]^ Change in the blood rheology occurs due to irregularities in the coagulation cascade that impact the vascular endothelial cells and cause clot formation.^[2]^ Thrombosis is a blood-related disease that occurs when veins or arteries get blocked due to the formation of blood clots (thrombi).^[3]^ The risks associated include stroke, heart attack, ischemic conditions, organ damage, etc., due to the occlusion of the formed clots.^[4]^ There are reports that indicate the adverse effects of vaccination of COVID inducing thrombotic thrombocytopenia, a rare condition of blood where thrombosis occurs despite low platelet count.^[5]^ A clinical study suggested effective treatment for vaccine-induced thrombocytopenia using argatroban and immunoglobulin (IVIG).^[6]^ However, these treatments are costly and have associated side effects.^[7]^

A class of antithrombotic drugs known as anticoagulants lowers the blood clotting risk.^[8]^ They perform either by dissolving existing clots or by preventing the formation of new ones. These include warfarin, heparin, and commercial anticoagulants like EDTA.^[9]^ Heparin activates the antithrombin, which inhibits thrombin to help form clots, and is widely used as an anticoagulant and an antithrombotic agent.^[10]^ Heparin, however, is inconvenient for long-term use because it requires frequent monitoring as the risk of bleeding is associated with it, along with the route of administration, preterm exhaustion, high cost, and toxicity.^[7a, 11]^ On the other hand, commercial anticoagulants such as EDTA act as a calcium chelating agent, resulting in the depletion of calcium in the blood, making them useful for blood collection and analysis. Notably, calcium plays a significant role as a cofactor in the blood coagulation mechanism, especially in those that regulate the function, stabilization, and activation of various clotting factors.^[12]^ However, it is hazardous for direct human administration due to its non-biodegradable nature and severe toxicity.^[13]^

To address these challenges, many approaches have been developed, but they lack in various terms, including high cost, low efficiency, risk of bleeding, shorter half-life, and restricted therapeutic window of medication. ^[14]^ Henceforth, novel technologies and strategies are needed to mitigate these problems. Nanomedicines have gained attention since they were introduced for their targeted action, remarkable physicochemical and biological properties, and reduced adverse effects.^[15]^ Here, we report the development of a novel inorganic nanocomplex, i.e., potassium ferric oxalate nanoparticles (KFeOx-NPs), that show anti-coagulation activity in mouse models and prevent thrombosis of human blood. Oxalate, a component of NPs, aids in chelating calcium, which affects the cascade of blood coagulation. Further, to address issues related to safety, *in vitro* and *in vivo* toxicity studies were performed in human blood, chicken egg, and mouse models, demonstrating the biocompatible nature of the nanoparticles.

**Scheme 1:**
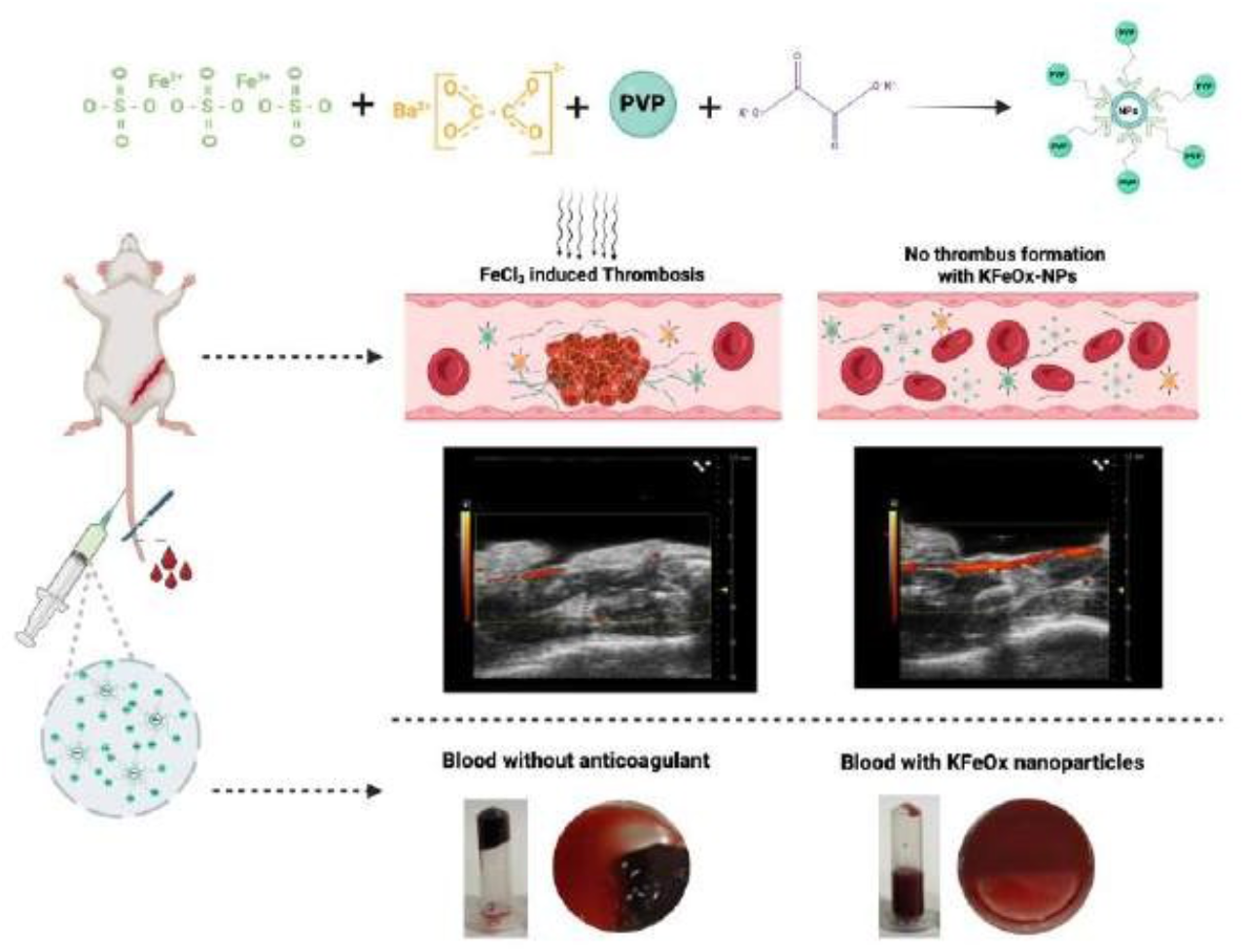
Schematic illustration of the KFeOx-NPs for anticoagulation and thrombosis prevention application

### 2. Results

### Structural analysis of synthesized nanoparticles

Oxalate is a natural calcium chelating agent by the formation of calcium oxalate.^[16]^ Calcium plays an important role in the blood coagulation pathways. A complex nanoparticle was developed to act as a calcium-chelating agent, preventing blood coagulation. PVP-stabilized potassium ferric oxalate nanoparticles (KFeOx-NPs) were synthesized by the simple inorganic complex reaction at room temperature by mixing ferric sulfate, barium oxalate, and potassium oxalate that was subjected to the steam bath for 3-4 hours, eventually changing the color from yellow-orange to lime green. After centrifugation, the colored solution precipitates out as nanoparticles of the inorganic complex denoted as KFeOx-NPs. A comprehensive characterization of the nanoparticles was performed (**Figure 1**). Dynamic Light Scattering (DLS) identified the size of KFeOx-NPs as 200–400 nm (**Figure 1a**) with a zeta potential value of -16.77; the negative value signifies stable nanoparticle production that is free from aggregation.^[17]^ X-ray diffraction spectroscopy (XRD) pattern displays distinct peaks at 2θ locations (**Figure 1b**). XRD analysis confirms the crystal structure of KFeOx-NPs supporting the SAED pattern (**Figure 1e**). **Figure 1c** displays the circular structure of KFeOx-NPs as seen through scanning electron microscopy (SEM) examination, which is confirmed in transmission electron microscopy (TEM) analysis (**Figure 1d**). This indicates that the size of NPs runs between 200 to 300 nm.^[18]^ The FTIR analysis of PVP (blue) and KFeOx-NPs (red) is illustrated in **Figure 1f**, with each sample demonstrating unique IR peaks. Significant differences in peak locations were identified in the FTIR spectra of KFeOx-NPs corresponding to PVP, which could be explained by the capping and stabilizing processes during NPs synthesis.^[19]^ A peak at 3435.08 implies -OH stretching, and it went on at 3443.27, showing PVP’s function in the synthesis and stability of KFeOx-NPs. Energy-dispersive X-ray analysis (EDAX) and color mapping from SEM confirm the presence of potassium (K), iron (Fe), carbon (C), and oxygen (O) elements in the synthesized NPs (**Figure 1g**). From the EDAX and ICPMS analysis, the formula of the synthesized NPs complex can be depicted as K_3_[Fe(C_2_O_4_)_3_]. The Brunauer-Emmett-Teller (BET) suggests a surface area of 2.9297 (m^2^/g) with a mean pore diameter of 0.5665 nm (**Table S1**). The reaction of NPs without using PVP was made as a control and characterized during the synthesis optimization. (**Figure S1-5**). However, these NPs were found to be less stable and possess larger sizes; hence, they were not used for any further studies. Following the synthesis, stability, and purity studies of PVP-stabilized KFeOx-NPs (denoted as KFeOx-NPs), they were employed for all the *in vivo* and *in vitro* biological studies.

**Figure 1:**
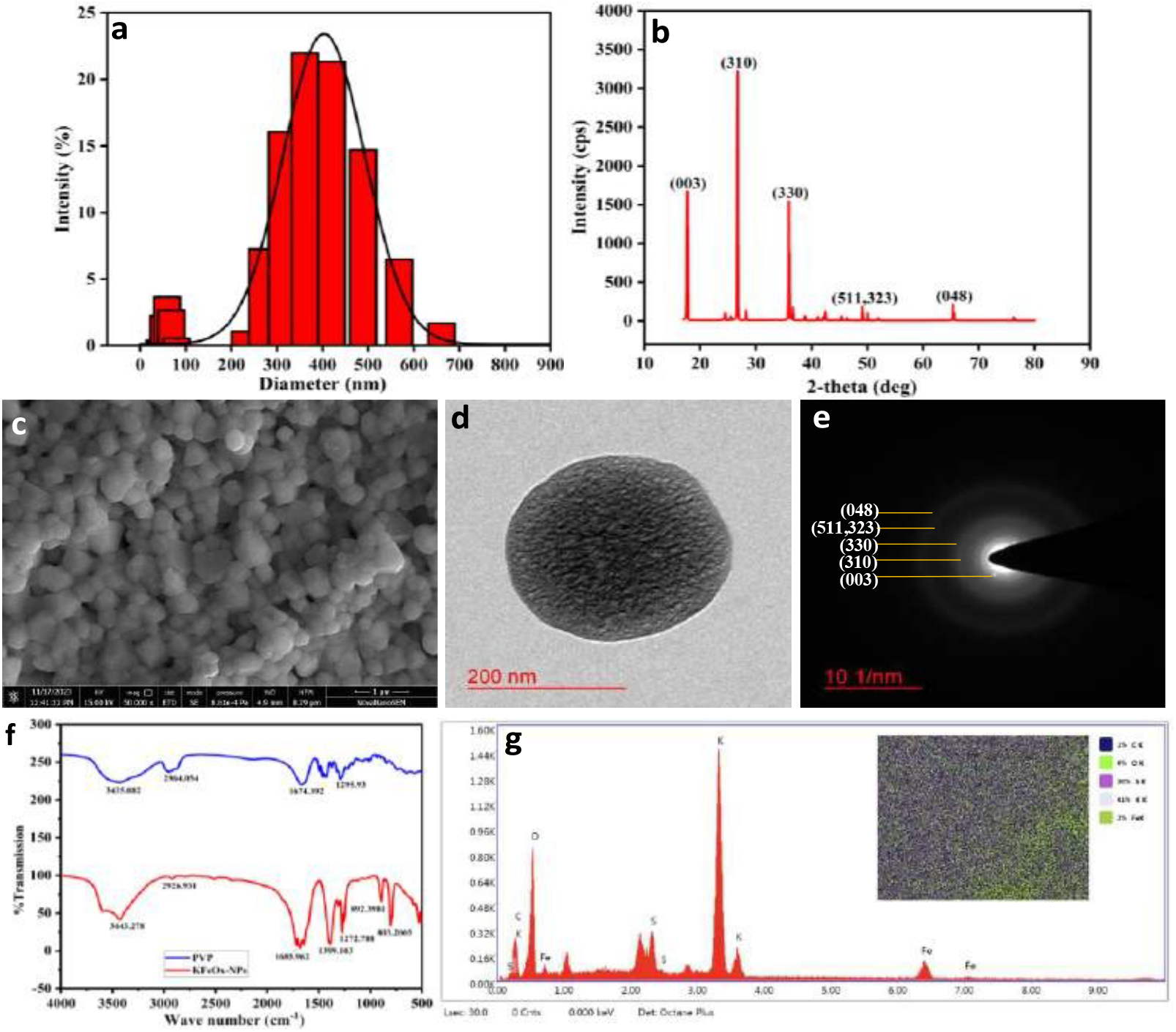
Characterization of KFeOx-NPs, (a) DLS showing size range of 200-400 nm, (b) XRD sharp peaks showing crystalline nature, (c) SEM images, (d) TEM image, (e) SAED pattern (f) FTIR of PVP (blue) and NPs (red), and (g) EDAX and color mapping.

### KFeOx nanoparticles prevent blood clotting

To confirm our hypothesis, anticoagulation studies of KFeOx-NPs were performed using human blood and compared with untreated and positive control, such as EDTA (**Figure 2a**). Visual observation suggests that NPs prevent the blood from clotting; a concentration of 2 mg/ml was selected for the *in vitro* assays. Commercial standards for determining clotting time are well-established and include prothrombin time (PT) and activated partial thromboplastin time (aPTT).^[20]^ For PT, blood containing KFeOx-NPs (2 mg/ml) takes longer to clot than commercial standard EDTA (**Figure 2b**). The typical range of PT in the blood is around 11 to 13 seconds. However, the presence of anticoagulant enables extension. As PT measures the clotting time when exposed to the tissue factor, which feeds back into the extrinsic pathway, the reagent used for the PT contains tissue factor (thromboplastin) and calcium that enables blood to clot. Hence, prolonged PT values suggest the effectiveness of NPs interfering with the extrinsic and common coagulation cascade, resulting in a delay in clot formation (**Video S1**).

**Figure 2:**
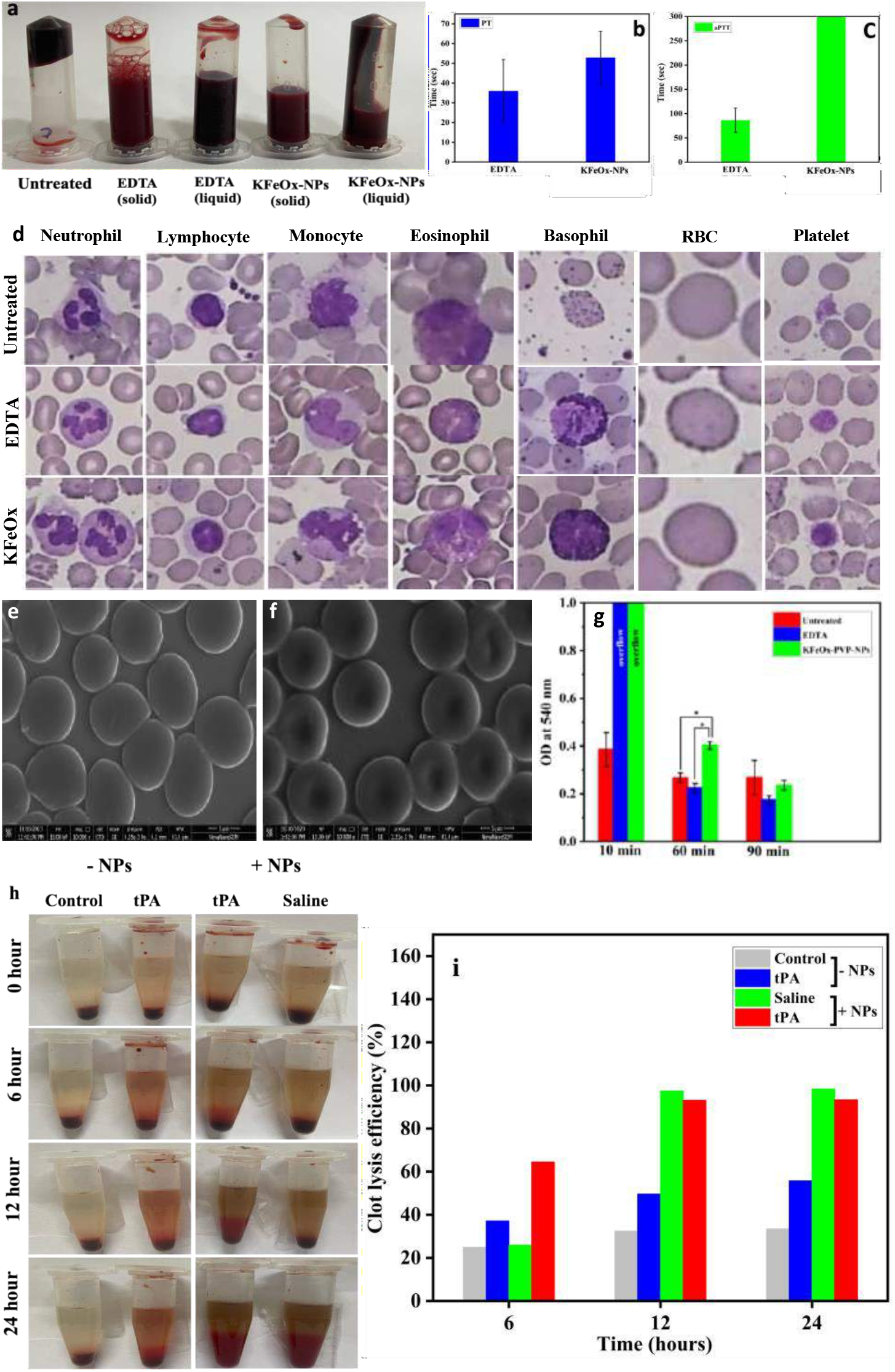
(a) Anticoagulating property of KFeOx-NPs, (b) Prothrombin time, (c) activated partial thromboplastin time, for blood containing EDTA and KFeOx-NPs (2 mg/ml), (d) high-resolution microscopical images of smears showing blood cells at 60x magnification, (e-f) SEM images of blood cell containing (e) EDTA, (f) KFeOx-NPs, (g) absorbance of dynamic clotting time assay at 540 nm, (h) clot lysis efficiency of pre-incubated KFeOx-NPs (2 mg/ml) in blood, with and without tPA (25 μg/mL), and (i) graph plotted from the clot lysis efficiency.

Conversely, blood containing KFeOx-NPs did not clot over 300 seconds (**Figure 2c**); the general range for aPTT in blood samples is typically around 25 to 35 seconds (**Video S2**). Prolonging aPTT values suggest effective interference with the intrinsic pathway of the coagulation cascade, which the aPTT primarily assesses.^[21]^ A microscopical analysis of blood with and without EDTA and KFeOx-NPs is shown in **Figure 2d**. Leishman stain was used to stain blood smears, which were then examined under a high-tech microscope (60x magnification). Blood components like red blood cells (RBC), white blood cells (WBC-basophile, eosinophile, monocyte, lymphocyte, neutrophile), and platelets were observed for all untreated, EDTA, and NPs containing blood showing no cellular morphological alterations in the blood. Additional studies, including morphological analysis using SEM (**Figure 2e-f**) and confocal images of blood smears, were performed, showing the intact size and shapes of blood cells following incubation with KFeOx-NPs, indicating blood compatibility **(Figure S6**).

The *in vitro* potentials of KFeOx-NPs dynamic clotting time assay were investigated to evaluate the anticoagulation property of dynamic clotting time. **Figure S7** shows the time-dependent clotting tendency of blood incubated with respective anticoagulants (EDTA and KFeOx-NPs) when induced with the calcium chloride solution. KFeOx-NPs show higher absorbance at each time point than EDTA and the untreated group (**Figure 2g**). The hydrolysis of RBC releases hemoglobin, whose absorbance is recorded at 540 nm.^[22]^ Hence, more absorbance demonstrates that more RBC concentrations in NPs exhibit longer clotting time in CaCl_2_-induced assay. To evaluate the thrombolytic potentials of KFeOx-NPs, the clot lysis efficiency was evaluated based on the weight of the thrombus over the period of time. This suggests that NPs possess no thrombolytic activity when treated to the existing blood clots (**Figure S14**). However, blood pre-incubated with the KFeOx-NPs is prone to form weaker clots in the presence of thrombin and has lysed over time with or without the addition of tPA (25 μg/mL) (**Figure 2h-i**). These results support the strong anti-thrombotic effects of KFeOx-NPs.

### Calcium saturation inverse the role of KFeOx-NPs in the blood

KFeOx-NPs were saturated with calcium (Ca) using CaCl2 to understand the molecular mechanism of blood clotting. Since literature suggests the ability of oxalates to chelate the calcium,^[23]^ the calcium saturation was performed at a ratio of 1.5 times and 3 times of oxalate. The structural change after the saturation of calcium was determined through the SEM and EDAX analysis (**Figure 3a-b**). This study shows the morphological changes in the NPs after adding the calcium. From EDAX analysis (**Figure 3b**), the amount of calcium bound to the KFeOx-NPs was determined as 4.2% and 6% for 1.5Ca and 3Ca, respectively. Hence, the molecular formula was determined as K_4_[Ca(C_2_O_4_)_3_] (denoted as K_4_CaOx_3_) and K_2_Ca[Ca(C_2_O_4_)_3_] (denoted as K_2_Ca_2_Ox_3_) for 1.5Ca and 3Ca; respectively. Blood coagulation was increased drastically with the incubation of K_4_CaOx_3_ and K_2_Ca_2_Ox_3_; the clotting time is plotted in **Figure 3c**. The results indicate that the K_2_Ca_2_Ox_3_ was more capable of clotting the blood (∼4 min) than the K_4_CaOx_3_ (∼10 min). Inset of **Figure 3c** further supports the observation.

**Figure 3:**
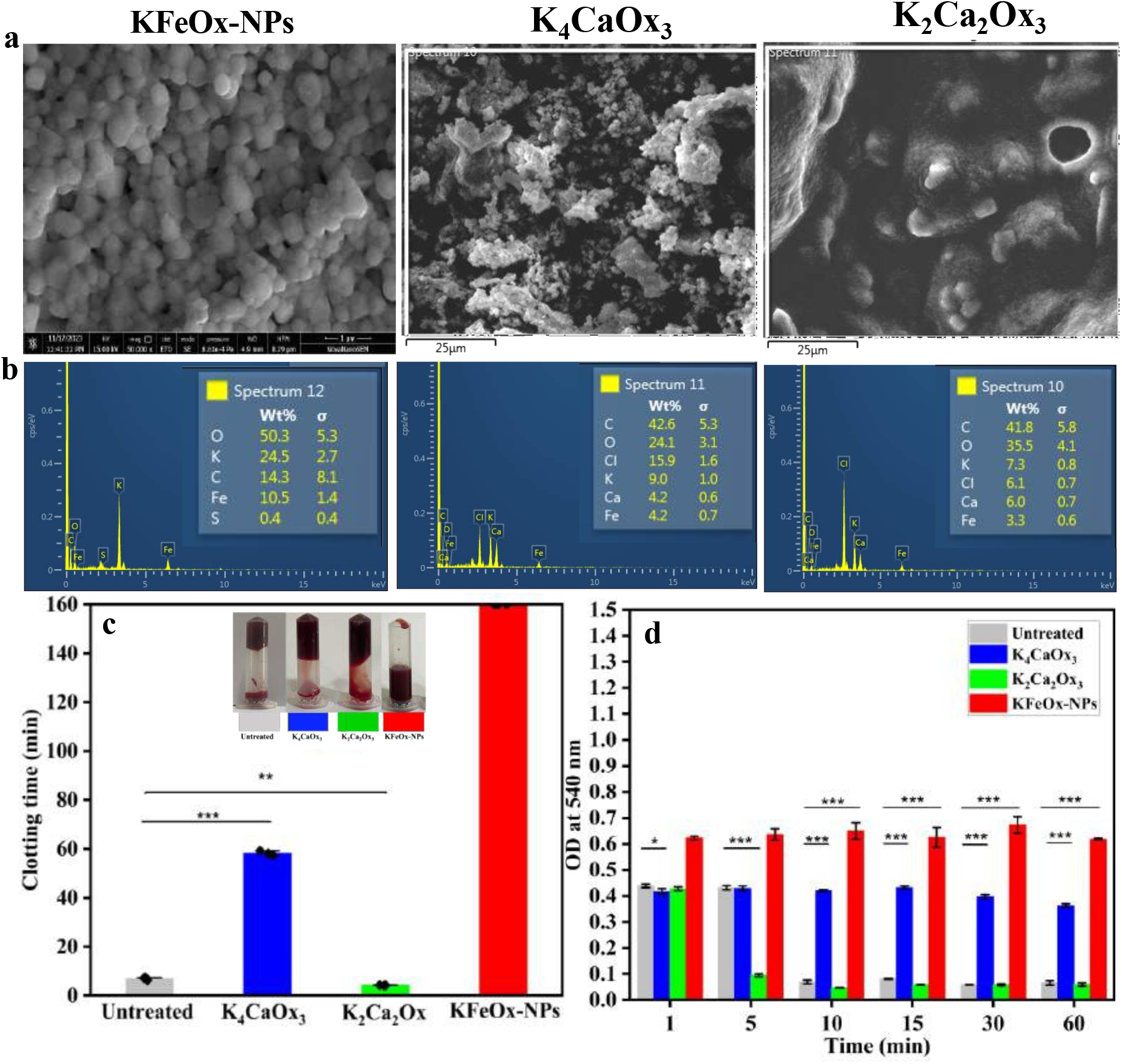
Calcium saturation study. (a) SEM images (b) EDAX of KFeOx-NPs, K_4_CaOx_3_ and K_2_Ca_2_Ox_3_, (c) clotting time of blood, and (d) absorbance of blood containing KFeOx-NPs, K_4_CaOx_3_, K_2_Ca_2_Ox_3_, and untreated.

These results suggest that the complete saturation of calcium to chelate the oxalates from the complex diminishes the anti-coagulation properties of the KFeOx-NPs. This confirms the blood clotting mechanism via the formation of a calcium-oxalate chelate complex, reducing the calcium concentration of the blood and preventing blood clots. Dynamic clotting time assay was further performed, confirming similar observations (**Figure 3d**). Calcium oxalate formation following the calcium saturation in the NPs was further confirmed using well know reaction of ammonia and ammonium oxalate.^[24]^ The formation of calcium oxalate precipitates the reaction, as shown in **Figure S8**, which further supports the molecular mechanism of blood clotting prevention of KFeOx-NPs.

### Toxicity analysis of KFeOx-NPs ensures biocompatibility

Influenced by the exceptional anticoagulating property, we next explored the *in vitro* biosafety of KFeOx-NPs in the human embryonic kidney cells (HEK-293 cell line) for 24 hours.^[1]^

The cell viability was analyzed after adding MTT reagent to the cells treated with different doses of KFeOx-NPs (1, 2, 4, 8 mg/ml) (**Figure 4a)**. Nearly 100% of the cells were found alive after the *in vitro* therapeutic dose of 2 mg/ml KFeOx-NPs, which ensures minimal cytotoxicity. However, with the increase in the concentration, the viability tends to decrease for 4 mg/ml (∼80% viable) and 8 mg/ml (∼40% viable). Following this, the blood compatibility of KFeOx-NPs was evaluated by the hemolysis assay shown in **Figure 4b**. The concentration at 2 and 4 mg/ml of KFeOx-NPs and the hemolysis rate were observed at 0.15% and 0.17%, respectively, indicating hemocompatibility of the KFeOx-NPs.^[25]^

**Figure 4:**
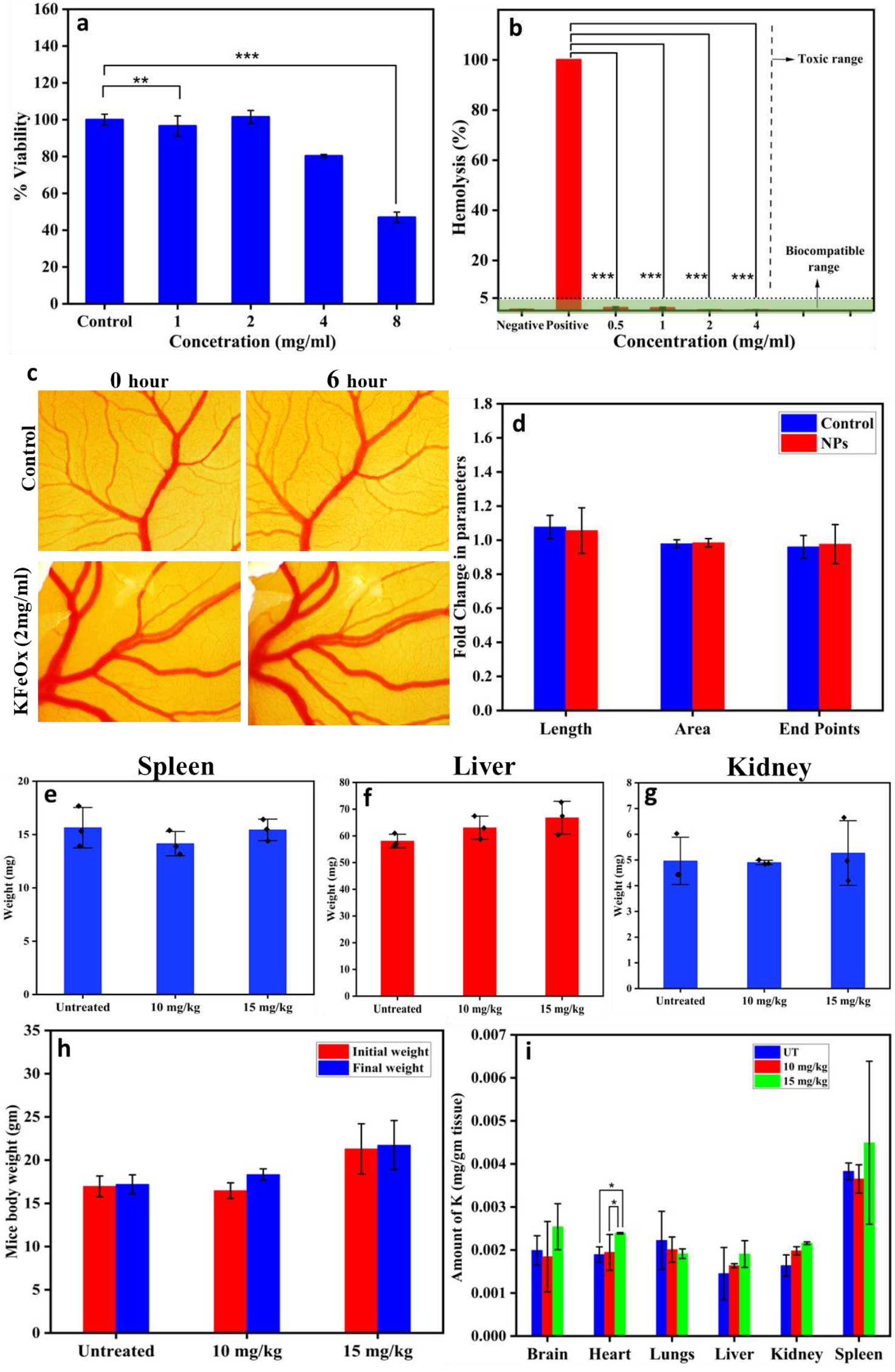
(a) Cell viability assay, (a) hemolysis assay, (c) Microscopic images of untreated and KFeOx-NPs treated infectious chick embryo models at 0 and 6 hours, (d) quantification with respect to length, area, and endpoints using ImageJ and Angiotool software, (e-g) weight of tissue normalized with body weight; (e) spleen, (f) liver, (g) kidney, (h) weight of animals before and after the acute toxicity study, (i) biodistribution of potassium by ICPMS analysis.

After that, we investigated the biocompatibility of the KFeOx-NPs *in ovo* in the chicken egg embryo model, known as the CAM assay.^[26]^ Chick embryos incubated with KFeOx-NPs for 6 hours showed the formation of mature blood vessels without any damage, including area, endpoints, and length, similar to the untreated embryos (**Figure 4c; Figure S9**). The investigation from **Figure 4d** clearly suggests no damage to the vascular network with the treatment of KFeOx-NPs at the dose of 2 mg/ml indicating *in ovo* biocompatibility.

Next, we tested the maximum safe dose (MSD) for toxicity assessment in the BALB/C mice. Seven days of toxicity study with the selected doses of 10 and 15 mg/kg of KFeOx-NPs suggests no symptoms of mortality and toxicity throughout the week.^[27]^ All the essential organs of mice (brain, heart, lungs, liver, kidney, and spleen) were harvested after one week (image captured shown in **Figure S10**) and weighed for the spleen, liver, and kidney shown in **Figure 4e-g** and other organs in **Figure S11**.^[28]^ All the animals in each group were observed to be healthy, with the absence of unusual behavior. Animals show significant weight gain in all groups (**Figure 4h)**, ensuring the nontoxic nature of the KFeOx-NPs.^[29]^ Following the toxicity studies, a 10 mg/kg dose was selected as a safe dose for all in vivo studies. Biodistribution studies were performed using ICPMS, which provides valuable insights regarding nanoparticle deposition in the vital organs (brain, heart, lungs, liver, kidney, and spleen).^[30]^ The investigation suggests maximum deposition of potassium (K) in the spleen for both doses (10 and 15 mg/kg) (**Figure 4i**).

Moreover, histopathological analysis using H&E staining also indicated the non-toxicity of the KFeOx-NPs on the tissue level. Histological imaging of all the vital organs after acute exposure to KFeOx-NPs (10 and 15 mg/kg) was shown in **Figure 5**. The pathological analysis indicates no sign of variation, like degeneration, inflammation, immune cell infiltration, or necrosis. Therefore, the nature of KFeOx-NPs at both 10 and 15 mg/kg doses can be considered non-toxic and useful for biological applications.

**Figure 5:**
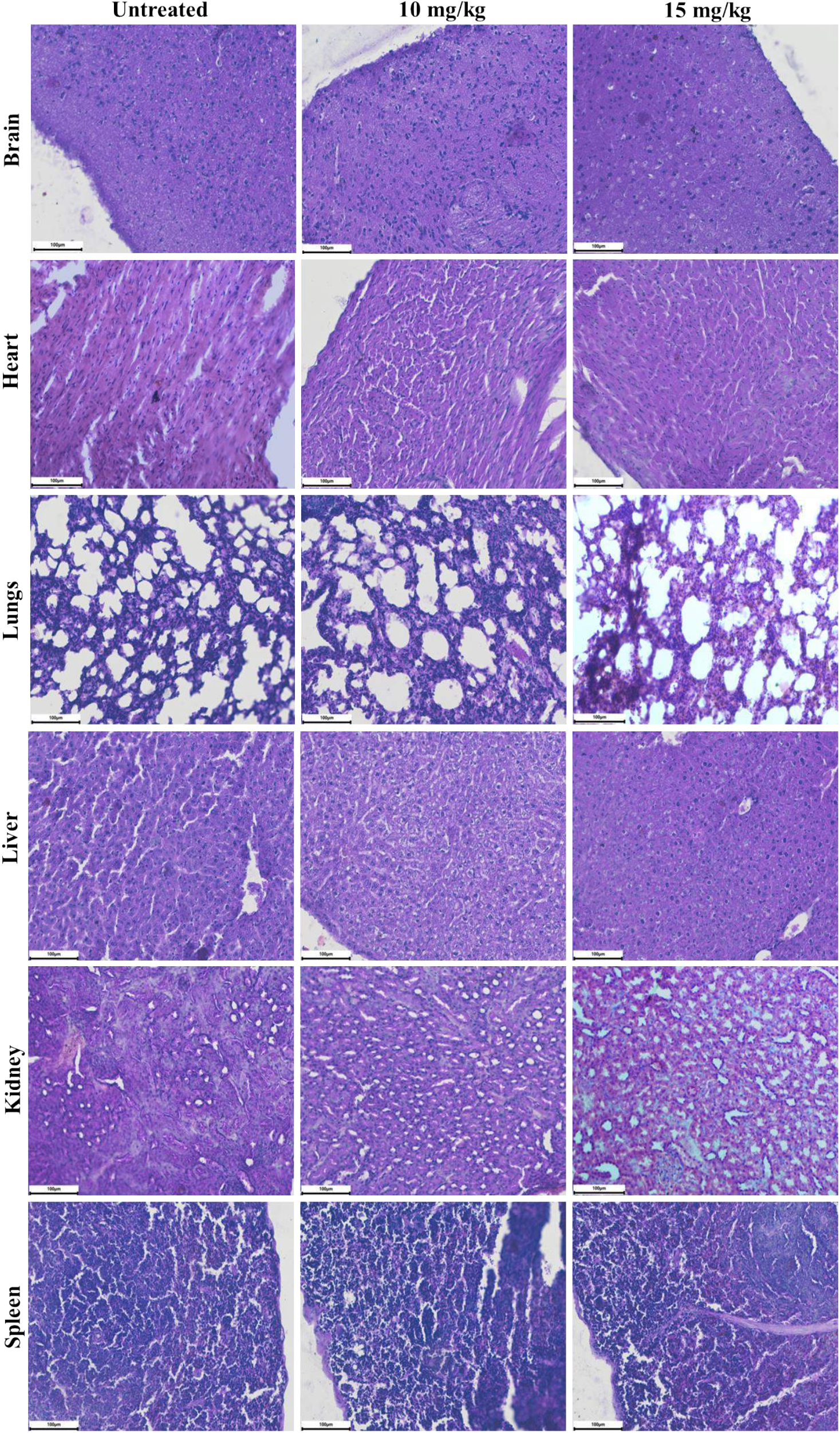
Histopathology images from essential organs (brain, heart, lungs, liver, kidney, and spleen) of BALB/C mice. Tissue sections of organs treated with KFeOx-NPs (10 and 15 mg/kg) and untreated were analyzed using H&E staining (n=3).

### KFeOx-NPs extend the clotting time and prevent thrombosis in a mouse model

The ability of KFeOx-NPs to show anticoagulation was evaluated in BALB/C mice (**Figure 6a**). Blood loss study from the mouse tail is an extensively used method to determine blood clot-related analysis.^[31]^ The KFeOx-NPs were injected intravenously (10 mg/kg) into the mice, and blood clotting time to stop leaking from their tails following the incision was recorded. The primary purpose of measuring animal bleeding time is to evaluate anti-clotting drugs’ hemorrhagic characteristics.^[32]^ Since prolonging clotting time indicates the delay in the clot formation process due to the addition of an anticoagulant in the blood, KFeOx-NPs show remarkably extended clotting time (**Figure 6b**). The group treated with heparin and KFeOx-NPs experienced nearly equal blood loss, higher than the untreated control group (**Figure 6c**). Furthermore, the absorbance of the blood collected from the tail of mice was measured (**Figure 6d**).

**Figure 6:**
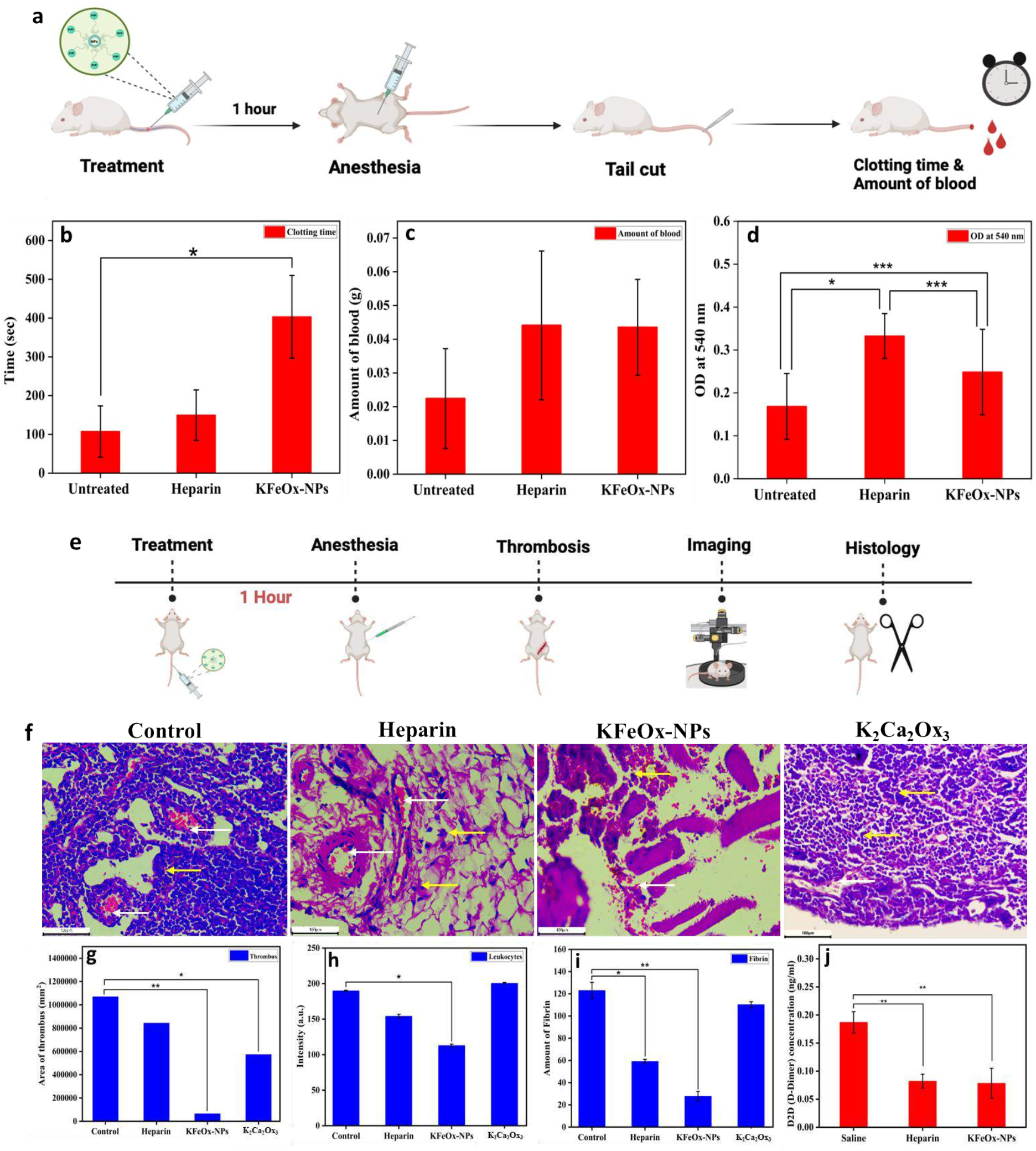
(a) Schematic representation of tail bleeding model, (b-d) parameters analyzed from in vivo assay including clotting time, amount of blood, and OD of blood at 540 nm, (n =3), (e) schematic representation and process of ferric chloride-induced thrombosis model, (f) histology images of the femoral vein, from the mice treated with saline, heparin, KFeOx-NPs, and K_2_Ca_2_Ox_3_, (g-i) histological parameters analyzed like area of thrombus, leukocytes, and fibrin using ImageJ software, and (j) D2D (D-Dimer) concentration estimated by ELISA test.

Encouraged by this, we then investigated the feasibility of KFeOx-NPs for thrombosis prevention. FeCl_3_ offers the basis for developing a ferric chloride-induced model as it causes oxidative damage to the vascular endothelial cells.^[33]^ Hence, this model is frequently used to investigate antithrombotic drugs and thrombogenesis (**Figure 6e**).^[34]^

The impact of the treatment is significantly influenced by systemic circulation, particularly in the case of blood vessels. Hence, one hour of treatment time was provided to facilitate the NPs circulation after the intravenous injection. After the experiment, the femoral vein was collected for section by sacrificing the animals. The embolism in the vein was then observed by staining H&E in vascular sections. Histology suggests the formation of a thrombus in the generated model, although the degree of thrombus was reduced following the treatment (**Figure 6f**). The microscopical histology analysis suggests a potent thrombosis inhibition effect in the femoral vein following KFeOx-NPs treatment. Based on the histological images, the analysis of the parameters involved in forming a thrombus was quantified. Accumulation of RBCs in the blood vessel is the first sign of thrombosis (denoted by white arrows) and was observed as minimal in the group treated with KFeOx-NPs (**Figure 6g**).

The presence of leukocytes indicates the involvement of immune reaction, here, due to vascular damage caused by the FeCl_3_ induction, observed less due to NPs (**Figure 6h**). Similarly, the presence of fibrin, as it exhibits to form a blood clot (**Figure 6i**), was quantified and found to be minimal in the KFeOx-NPs treated group. Interestingly, the K_2_Ca_2_Ox_3_ treated group revered the thrombosis-preventive effects of the KFeOx-NPs, confirming that calcium chelation to oxalates plays a crucial role in preventing blood coagulation and thrombosis.

The concentration of mouse D-dimer, a biomarker derived from the breakdown of fibrinogen in the blood clot dissolution process, was measured from serum separated from mouse blood. A higher level of D-dimer concentration was observed in the control sample treated with saline, suggesting more formation of thrombosis than in heparin and KFeOx-NPs treated groups (**Figure 6j**). This supports the thrombotic prevention effects of KFeOx-NPs.

Power Doppler and ultrasound imaging were applied before and after the treatment to track the blood flow (**Figure 7a-b**). Power Doppler and flow Doppler images clearly show the typical blood flow in the femoral vein of all the groups before thrombus induction.^[35]^ After the thrombus induction with the topical application of FeCl_3_-soaked filter paper on the vein, imaging was performed to monitor the effect of various treatments in thrombus formation (**Figure 7a-b, Figure S12**).^[36]^ Impressively, the KFeOx-NPs treated group prevented the thrombus formation after FeCl_3_ induction, as validated by the uninterrupted blood flow and mean velocity calculated from the flow doppler (**Figure 7a-c**).^[37]^ Power Doppler images confirmed the blood clumping in the femoral vein and interrupted blood flow for untreated control, heparin-treated group, and K_2_Ca_2_Ox_3_-treated groups, indicating thrombus formation (**Figure 7b)**. These significant changes inside the vein before and after the treatment clearly confirm the thrombosis prevention property of KFeOx-NPs (**Figure 7c**).^[38]^ Importantly, the treatment of K_2_Ca_2_Ox_3_ (10 mg/kg) intravenously in the mice demonstrated the formation of thrombus and reduction of blood mean velocity similar to that of the control group (**Figure 7c)**. This further supports the hypothesis that KFeOx-NPs act as a calcium scavenger by creating a calcium-oxalate chelation complex that prevents blood coagulation and thrombosis.

**Figure 7:**
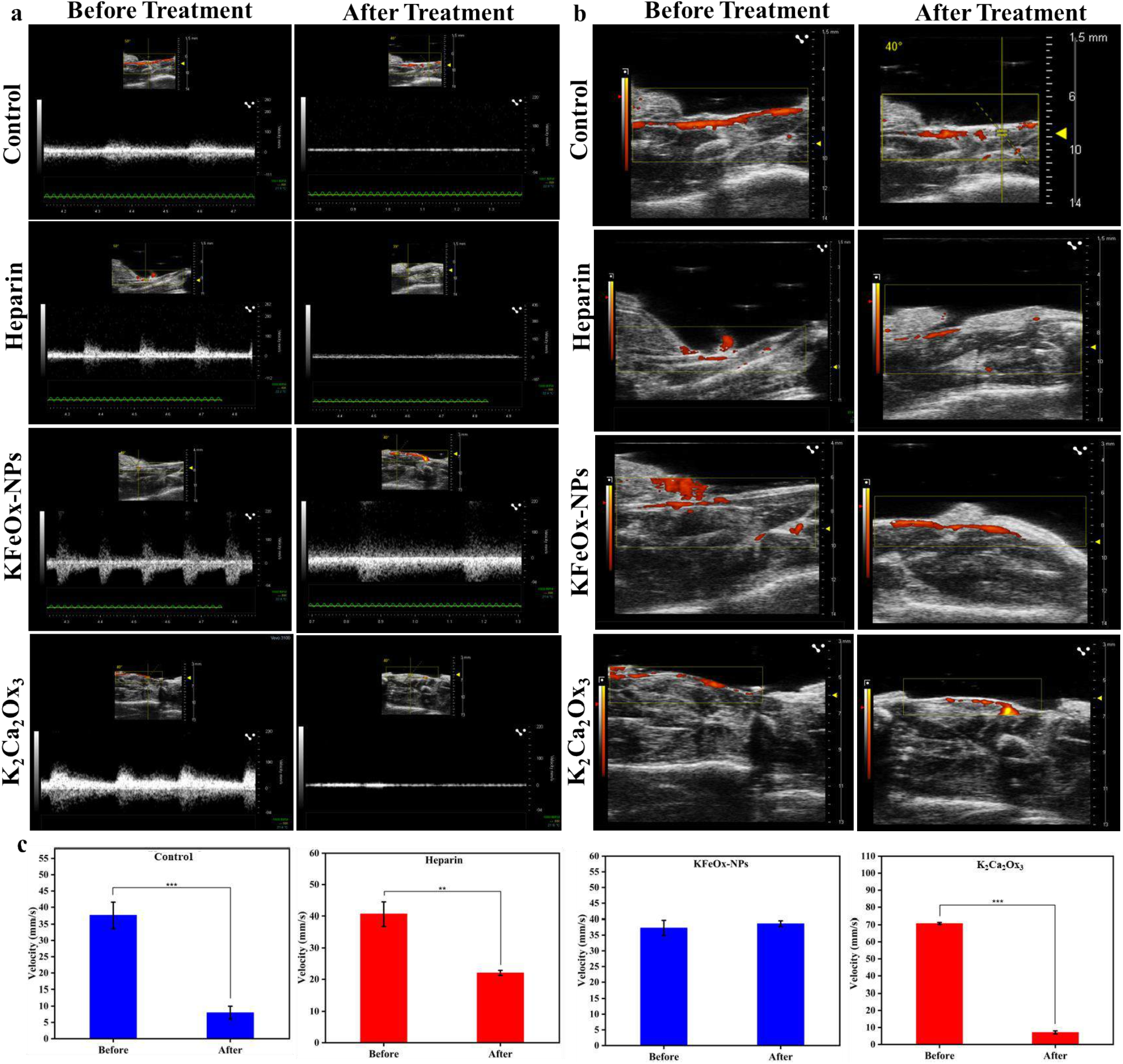
In vivo thrombosis imaging in BALB/C mice analyzed by Vevo software. (a) Ultrasound images showing the blood flow in the femoral vein before and after the treatment, (b) power doppler images showing the presence of blood in the femoral vein before and after the treatment of saline, heparin, KFeOx-NPs, and K_2_Ca_2_Ox_3_, and (c) mean velocity plot of all the groups calculated from the flow doppler images of the vein before and after the treatment (n=6).

### KFeOx-NPs prevent blood coagulation and protein attachment when coated with a catheter

To test the effectivity of NPs against the components of blood concerning medical devices, the silicon catheters were plasma treated and then coated by dipping in the saturated solution of KFeOx-NPs and EDTA (**Figure 8a-b**). After 4 hours of incubation with human blood, SEM was used to analyze the surface properties of the untreated and treated catheters (**Figure 8a**), which confirms the coating and presence of cellular components of the blood. Uncoated and EDTA-coated catheters show the presence of RBC (white arrows) on the surface, whereas NP-coated catheters show the absence of blood cells over the surface. This result indicates that NPs coating is capable of prohibiting the formation of blood clots when used to coat medical devices or implants.^[39]^ In addition to that, the presence of attached protein was estimated using CBB staining. The result suggests that the surface of the catheters coated with KFeOx-NPs shows less attachment of the blood protein when compared to EDTA and untreated catheters (**Figure 8c**). KFeOx-NPs prevent blood coagulation inside the catheter and reduce protein attachment.^[40]^ Moreover, the flow properties of these blood-incubated catheters (uncoated and KFeOx-NPs coated) were measured using a syringe pump by measuring the time taken to pass 5 mL of 10% serum solution (**Figure S13**). The result suggests that the KFeOx-NPs-coated catheter took the least time when compared to uncoated and EDTA-coated catheters (**Figure 8d**). The catheter with just ∼1.5 cm in length showed a difference in the flow time, which can be multiplied by increasing the catheter length. A typical size of central venous catheters used for long-term infusion therapies is 15-20 cm.^[41]^ Hence, a significant reduction of the flow time can be obtained with the catheters coated with KFeOx-NPs, improving their functions and applications. This is particularly crucial in medical settings where catheters must remain unclogged for extended periods.

**Figure 8:**
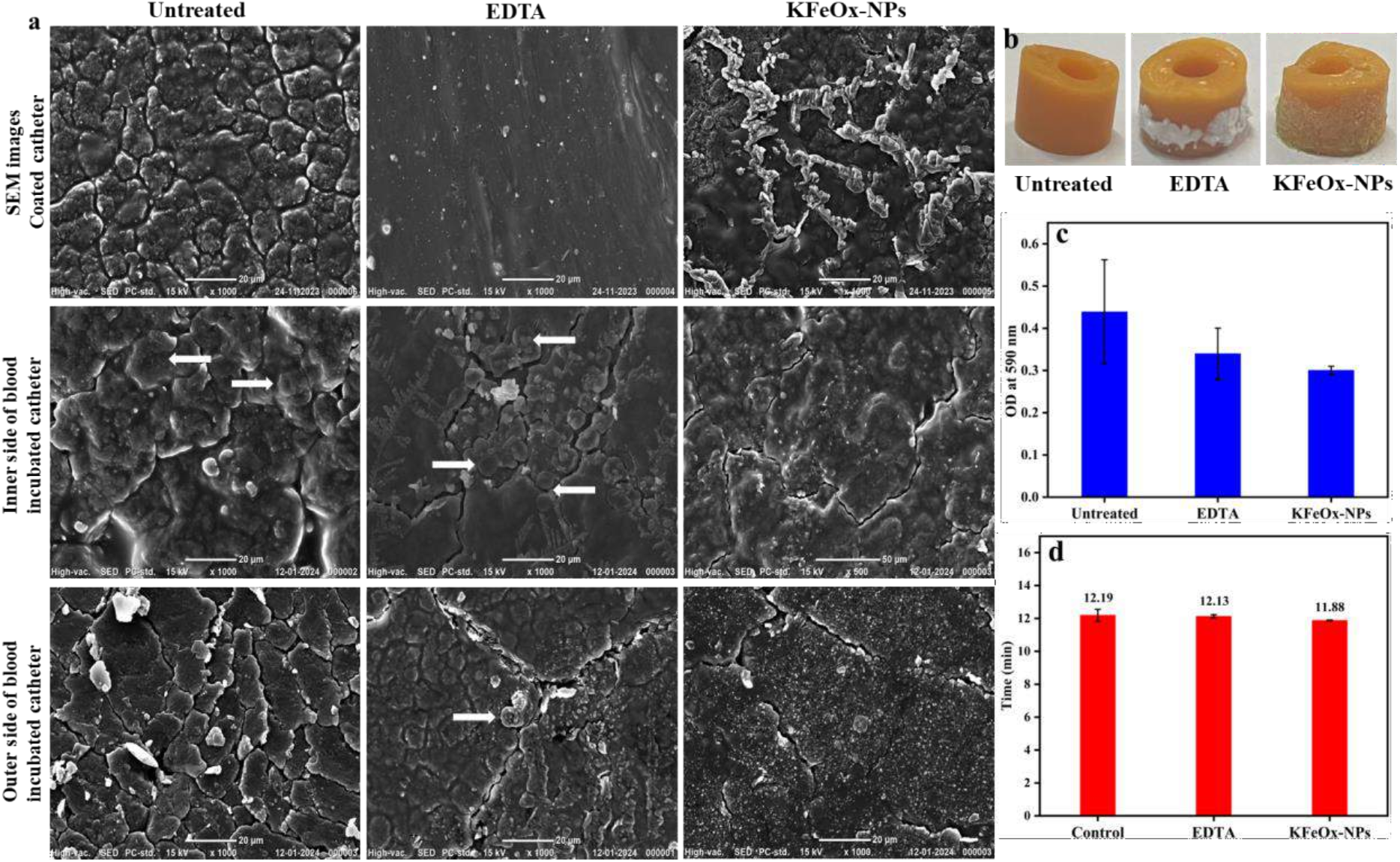
Application of NPs using a catheter. (a) SEM images of the catheter before and after the coating of EDTA and KFeOx-NPs, (b) images of the coated catheters, (c) protein estimation from the catheter after incubating with blood, and (d) flow ability of catheters tested after blood incubation.

## 3. Discussion

We reported the synthesis of novel PVP-stabilized potassium ferric oxalate nanoparticles (KFeOx-NPs) for blood clot management. KFeOx-NPs incubated with human blood can inhibit the blood from clotting with a minimum dose of 2 mg/ml and further prevent thrombosis in an animal model at 10mg/kg dose. Additional studies demonstrated that blood pre-incubated with KFeOx-NPs is prone to lysis due to the formation of weak clots.

The KFeOx-NPs are readily soluble in water, which aids their easy absorption and distribution throughout the body and speeds up their onset of action. In addition, their behavior is often more predictable than that of lipid-soluble drugs, which can accumulate in fat tissues and have variable release patterns. To further analyze the mechanism associated with the activity of KFeOx-NPs, in compliance with the existing research, we examined the calcium chelation mechanism of oxalate by saturating KFeOx-NPs with CaCl_2_.^[42]^ This demonstrated that KFeOx-NPs efficiently chelate the calcium ions from the blood and alter the coagulation cascade, preventing fibrin from forming clots. Since calcium is necessary in blood coagulation, to be an effective anticoagulant, chelation of calcium is one efficient way to prohibit the activation of the factors that cause fibrin production. There are reports that use a similar mechanism for anticoagulation.^[43]^ However, KFeOx-NPs are low in cost, highly biocompatible, and water-soluble, resulting in improved applicability in blood clot management.

The cascade of blood coagulation involves two major pathways: intrinsic and extrinsic. The common pathway of coagulation is then generated by converging both pathways, after which a series of events are led by the activated Factor X. The inactive enzyme prothrombin (Factor II) is converted to its active form, thrombin (Factor IIa), by prothrombinase. Thrombin then transforms soluble fibrinogen (Factor I) into insoluble fibrin strands. With the support of Factor XII, these fibrin strands lead to blood clot formation. Notably, calcium plays a significant role as a cofactor in the blood coagulation mechanism, especially in those that regulate the function, stabilization, and activation of various clotting factors.^[12]^ The commercial standard clotting time assays, PT and aPTT, also clarified the precise mechanism of coagulation obstruction by KFeOx-NPs. Unlike PT, aPTT continued without clotting, validating the KFeOx-NPs interference in the intrinsic pathway where it functions. The main components of the aPTT reagent that act as contact/surface activators are ellagic acid, kaolin, and silica, which all influence factor XII activation. A kaolin clotting time assay was used to recognize the activation of factor XII after incubation and demonstrate the intrinsic pathway at which the KFeOx-NPs alter the coagulation process (**Video S5)**. After kaolin was added, the test progressed without clotting the plasma, indicating that the KFeOx-NPs restrict the activation of factor XII, which corrupts the coagulation cascade and prevents other factors from activating to create fibrin clots (**Video S5)**.^[44]^ In contrast to many other clotting factors, limiting Factor XII often does not raise the risk of bleeding, which makes Factor XII inhibitors potentially valuable as anticoagulants.^[45]^ An anticoagulant can efficiently restrict the start and spread of blood coagulation through various routes when it possesses the capacity to both chelate calcium and block Factor XII activation. This dual mechanism guarantees a strong anticoagulation effect, making it appropriate for various medical and research applications, including anti-coagulation, clot management in medical devices, and thrombus prevention.

**Scheme 2:**
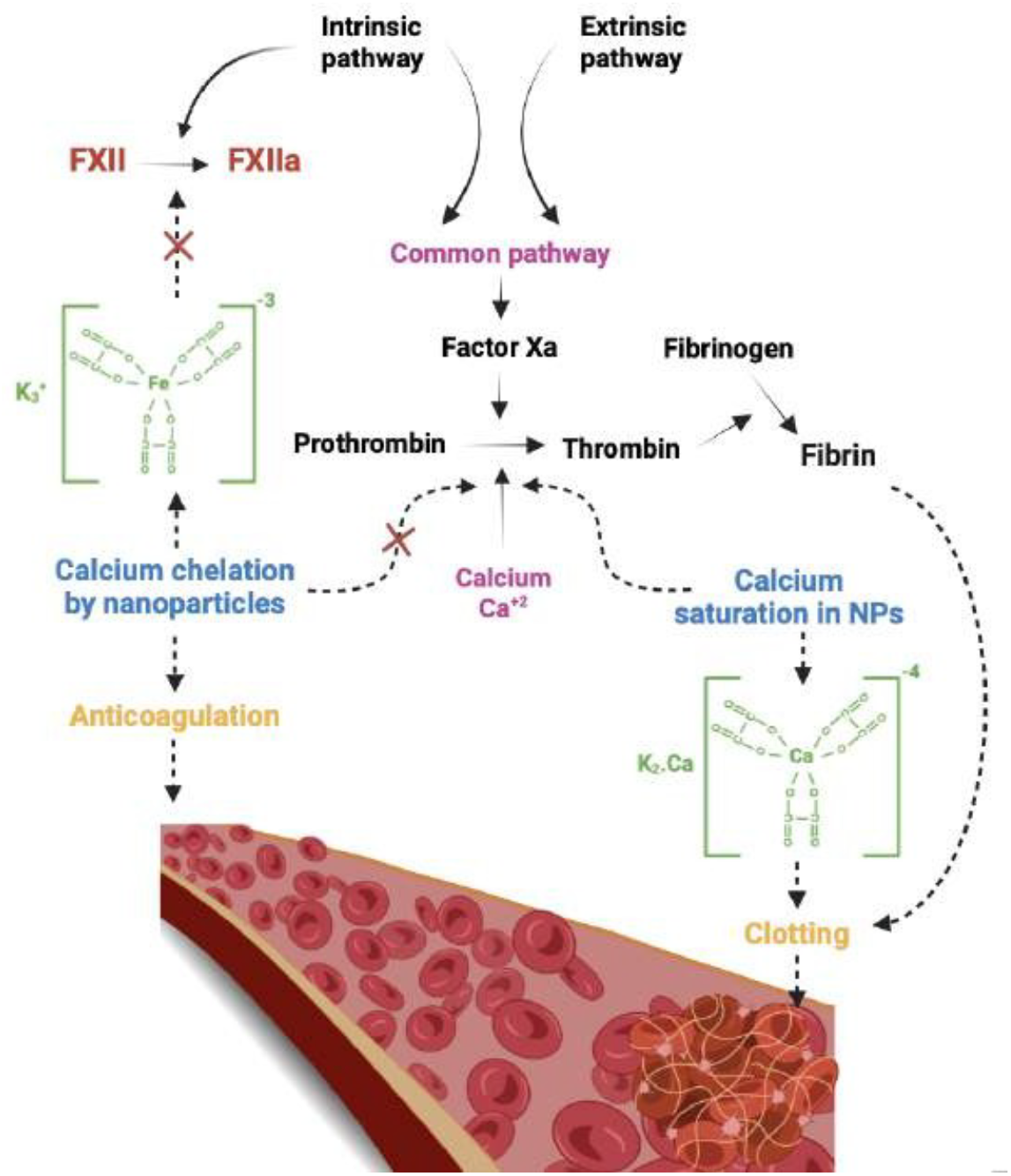
Representation of the mechanism of the KFeOx-NPs.

Other studies suggest that the anticoagulant formulation relied on the RNA-DNA fibers, which increases the complexity of synthesis and purification, leading to an increase in the platform’s overall cost and production time.^[10]^ On the other hand, our NPs are synthesized through simple inorganic complex reactions, which is an inexpensive and rapid process. There are studies that report the surface modification of the nanomaterial to enhance the biological effect in the system; for instance, *Zhou et al*., mentioned the surface modification with the low molecular weight heparin as a shell on the nanocomplex, which may increase the complexity of the formulation, potentially impacting the reproducibility and scalability of the manufacturing process.^[9]^ Compared to other studies, most nanoparticle-based therapies have used nanomaterial as a delivery vehicle loaded with antithrombotic drugs. A study by *Li et al*., explained the targeted delivery of hirudin biomimetic nanoplatelets for thrombosis prevention.^[14]^ Similarly, plasminogen activators were delivered through engineered nanoplatelets by Xu et al., in multiple mouse thrombosis models.^[28]^ In this work, the KFeOx-NPs act as a therapeutic agent without any functional modification. *Lu et al*., conducted dose-dependent PT and aPTT with heparin-like anticoagulant polypeptides, demonstrating the clotting time values, which are lower than our KFeOx-NPs.^[46]^ Similarly, coumarin-derived anticoagulants show an inhibitory effect on platelet aggregation but possess lesser PT values than our NPs.^[47]^ Considering the above literature, KFeOx-NPs are low in cost, biocompatible in nature, and highly effective in preventing blood coagulation and thrombosis. Additionally, blood clots in the presence of KFeOx-NPs are weaker in nature and prone to lysis automatically in the presence of existing tPA present in the body.

KFeOx-NPs were used to coat the medical urinary catheter to reduce the risk of the formation of blood clots and protein attachment and improve the blood flow through them. In general, clot formation is a major concern with implants, as they may trigger nonspecific protein attachment, resulting in inflammation, fibrosis, and risk of infection.^[40]^ Coating with the anticoagulant might reduce the risk of blood clots and thrombosis, especially for blood-contacting medical devices like cardiovascular implants.^[41]^ For instance, *Wang et al*., demonstrated an effective functional coating strategy to resist coagulation as well as inflammation on cardiovascular devices.^[1]^ Similarly, KFeOx-NPs can be used to improve the durability of these medical devices, including cardiovascular implants and catheters.

The limitation of our study is that KFeOx-NPs do not exhibit thrombolysis activity on pre-existing blood clots. To solve this, our ongoing work is to combine the KFeOx-NPs with a lower dosage of the potent thrombolytic drug, for example, tissue plasminogen activator (tPA), to actively lyse the pre-existing clots. Moreover, research indicates that internal calcium chelation might result in hypocalcemia, a disorder that our initial toxicology studies have not shown in BALB/C animal models. However, detailed studies are required to confirm the long-term safety of the KFeOx-NPs for potential clinical use in humans for various blood clot management applications.

## 4. Conclusion

We reported the synthesis of the novel PVP-stabilized potassium ferric oxalate nanoparticles for blood clot management. KFeOx-NPs act as anticoagulants in human blood and can be used for thrombosis prevention in a mouse model. Further, we employed NPs to coat medical catheters, demonstrating the prevention of blood clots and reduced protein attachment, improving the blood flow properties. Overall, the potential application of this material holds great promise in biomedical applications, especially for blood clot-related diseases and medical implants like blood-contacting devices.

## 5. Experimental Section/Methods

### Materials

Ferric sulfate, Potassium oxalate, Polyvinyl pyrrolidone (PVP), Ethanol, EDTA, Heparin, Leishman stain, calcium chloride, Nutrient agar, kaolin, PBS, and MTT reagent were purchased from Sisco Research Laboratories Pvt. Ltd. (SRL) India. Barium oxalate was purchased from Otto Chemicals Pvt. Ltd., Dulbecco’s Modified Eagle Medium (DMEM), nitric acid from Finar Chemicals, and trypsin was purchased from HiMedia Laboratories.

### Cells and Animals

The Human embryonic kidney cells (HEK-293 cell line) were purchased from the National Centre for Cell Science (NCCS) Pune. The cells were cultured using DMEM media at 37 °C in a humidified atmosphere with CO_2_ (5%). Both male and female BALB/C mice (20-30 gm) were purchased from CDRI Lucknow. The mice were provided with limitless access to water and standard food. They were randomly selected for the experimental groups. Following a clearance of “the anticoagulating potential of nanomaterial/biomaterials for blood clot management” by the Institutional Animals Ethics Committee (IAEC) of Indian Institute of Technology, BHU, Varanasi approval ID: IIT(BHU)/IAEC/2023/II/075 has carried out all *in vivo* investigations.

### Synthesis of KFeOx Nanoparticles

The inorganic complex nanoparticles (KFeOx-NPs) were synthesized using the bottom-up technique. In a beaker (100 ml), 2.5 gm of ferric sulfate and 5 gm of barium oxalate, along with 250 mg of PVP (polyvinyl pyruvate) as a stabilizing agent, were added and mixed with 60 ml mili-Q water. This was stirred on the magnetic stirrer for 30 mins with 400-500 rpm at room temperature. 2.7 gm of potassium oxalate was added and stirred again for 15 mins. 4 ml of HCl (N/10) was added as a solubilizing agent to this mixture and continued stirring for 15-20 min. This reaction was transferred in a steam bath for 3-4 hours with occasional stirring. This compound was then filtered through filter paper and transferred equally into four centrifuged tubes with 15 ml of ethanol for centrifugation at 25 °C and 6500 rpm for 45 min until the color changed and a light green colored precipitate was obtained at the bottom. The supernatant was discarded, and the precipitate was washed twice (30 mins cycle each) with ethanol in the same centrifuged condition. The precipitate was air-dried after the removal of the supernatant. This precipitate or NPs were then collected and used for all characterization and biological studies.

### Ex vivo anticoagulation Activity

To determine the anticoagulation property of KFeOx-NPs, Prothrombin time (PT) and activated partial thromboplastin time (aPTT) were measured. Blood samples were collected from healthy volunteers; the protocol was approved by the ethical committee of the Institute of Medical Sciences, Banaras Hindu University (Ref. No. ECR/526/Inst./UP/2014/RR-20 date-May 19, 2020). The blood samples were collected in vials containing anticoagulants (EDTA and KFeOx-NPs). Platelet-rich plasma (PRP) was obtained after centrifugation of blood samples at 1500 rpm for 15 min. The PT reagent was pre-warmed before the test started, and the incubation of 100 μl of plasma, which was dispensed in a fresh cuvette containing a dispenser stirrer ball for 180 sec. 200 μl of PT was then added, and the timer was started. Similarly, in the APTT assay, CaCl_2_ and APTT were pre-warmed. 100 μl of plasma with 100 μl of APTT reagent was incubated for 180 sec. Further, 100 μl of CaCl_2_ was added, and the timer was started. EDTA is a commercially used anticoagulant and hence kept as control. PT and APTT were recorded (Thrombostat 1 Behnk Elektronik) in seconds.

To determine the thrombolytic activity of KFeOx-NPs, the 2 mg/ml dose of NPs was used. Initially, 100 μl of NPs incubated blood was added with 1 unit of thrombin to form a thrombus. The samples were then further incubated at 37°C for 3-4 hours followed by overnight incubation in 4°C. The samples were then incubated with either saline or tPA (25 μg) at 37°C and the weight of the thrombus was taken in various time points. The clot lysis efficiency of the thrombus was calculated as:

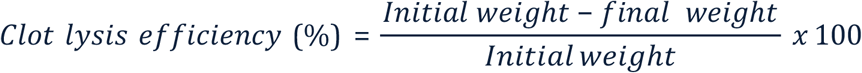

### In vitro anticoagulation assay

For the dynamic clotting time assay, 100 μl of blood (untreated, with EDTA, and with KFeOx-NPs) was added to a 6-well plate. Immediately, 0.2 M CaCl_2_ 10 μl solution was added, and incubated for 10, 60, and 90 mins. The samples were washed with MiliQ water for 1 minute, and the absorbance of the washed solution was detected at 540 nm in the microplate reader (Synergy H1-Microplate reader).

### Cell viability assay

MTT (3-(4, 5-dimethyl thiazolyl-2)-2, 5-diphenyltetrazolium bromide) assay is a colorimetric method used to evaluate the cytotoxicity of any substance. When mitochondrial dehydrogenase enzymes are active, this reagent forms formazan crystals, which turn purple from yellow by adding DMSO. Human embryonic kidney cells (HEK-293 cell line) (10,000 cells/well) were incubated for 24 hours in 96 well plates. The cultured cells were treated with different concentrations of KFeOx-NPs (0.5, 1, 2, 4, and 8 mg/ml) with further incubation of 24 hours. 100 μl of MTT reagent in DMEM (0.5 mg/ml) was then added to each well by replacing the previous media. After 4 hours of MTT, DMSO was added to solubilize the formazan crystals. The purple colour change was measured in the microplate reader (Synergy H1-Microplate reader) at 570 nm. The viability was calculated as:

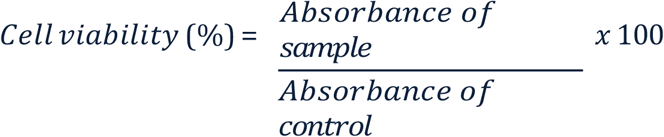

### Haemolysis assay

To test the KFeOx-NPs hemocompatibility, fresh human blood (4 ml) was collected and centrifuged for 15 minutes at 3000 rpm. Phosphate buffer saline (PBS) was used three times to wash the settled red blood cells (RBC). 50 μl of separated RBC was added to 900 μl of PBS, and the remaining 50 μl of the sample were treated at various doses (0.5, 1, 2, 4, and 8 mg/ml of KFeOx-NPs). These were then incubated for four hours while shaking. The optical density (OD) of the supernatant after centrifugation was measured at 540 nm in the microplate reader (Synergy H1-Microplate reader). PBS and triton X-100 were used as negative and positive controls, respectively. The percentage of erythrocyte lysed was calculated by the formula:

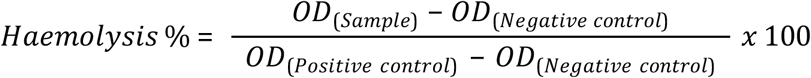

### Biocompatibility assay

The Chorioallantoic Membrane (CAM) assay is an extensively researched *in ovo* assay to assess biomaterials before being further examined in animal models. This assay was performed to examine the impact of KFeOx-NPs on the development of blood vessels. Fertilized eggs were first acquired, and then these were incubated for 4 days at 37°C. After the incubation phase, a small window was created on the eggshell to monitor the growth of the developing embryo. 6 mm Whatman filter paper (discs) was soaked with a mixture containing KFeOx (2 mg/ml) for a few seconds. These discs were subsequently placed above the CAM layer. At this early time, the images were captured employing a Mgnus MagZoom TZM6 Trinocular Stereo Zoom Microscope. The eggs were then sealed with the help of parafilm and placed back in the incubator for another six hours. Additional images were taken after 6 hours and 24 hours to measure the fold change in different parameters. As a point of comparison, untreated eggs were utilized. Quantification of the microscopic images was done using Angiotool and ImageJ software.

### Acute toxicity and biodistribution studies

Toxicity studies were performed according to the Organization for Economic Co-operation and Development (OECD) guideline no. 423 to investigate the therapeutic as well as toxic doses of KFeOx-NPs. For this study, female BALB/C mice were used to determine the maximum torrent dose (MTD). Mice were categorized into 3 groups, namely untreated, 10 mg/kg, and 15 mg/kg, as 3 mice per dose (n=3). The stock solution of 2 mg/ml in saline was used for the study. 200 μl of the dose, calculated according to the body weight of the mice, was prepared from the stock solution and administered intravenously. The observations, including behaviour, general health, and mortality, were noted on a daily basis for 7 days after the treatment. The mice in each group were weighed daily till they were sacrificed. The mice were sacrificed on the 8^th^ day of the treatment, and all vital organs (brain, heart, lungs, liver, kidney, and spleen) were harvested and fixed in formalin solution. The fixed organs were then sent for tissue sectioning in pathology labs, and histological analysis was done using a microscope. For another group, to exhibit biodistribution studies, the harvested mice organs were digested using nitric acid for 4-5 days. After the digestion, the solution was diluted with PBS made to the volume of 5 ml. This solution was then submitted for ICPMS after filtration through a 0.22 μm syringe filter.

### In vivo mouse tail bleeding model

The mouse tail-cutting bleeding model was performed to evaluate the anticoagulation potential of KFeOx-NPs. BALB/C mice (20-30 gm) were randomly selected and divided into 3 groups, namely untreated, heparin, and KFeOx-NPs treated (n = 3). These were intravenously injected with 200 μl of 1 mg/ml solution of heparin and NPs in saline, irrespective of their body weight. The animals were anesthetized with ketamine (60 mg/kg) and xylazine (7 mg/kg) after one hour of treatment, and the tail was amputated (3-4 cm above the tip). Parameters, including clotting time and amount of blood, were noted as soon as the tail cut. The optical density of obtained blood was measured by adding 10 ml Mili-Q to the blood at 540 nm in the microplate reader (Synergy H1-Microplate reader).

### FeCl_3_-induced thrombus model

The ferric chloride-induced model is most widely used for thrombosis investigations The thrombus model was constructed by anesthetizing BALB/C mice. The incision was made near the lower limb to expose the femoral vein. The fat and connective tissue layers were removed carefully to isolate the vein. Filter paper soaked in 10% FeCl_3_ solution was placed over the exposed vein for 3-5 mins to induce thrombosis.

### In vivo thrombus imaging by Ultrasound and Doppler flow

The BALB/C mice (20-30 gm) were randomly selected and divided into 3 groups (n = 6). The animals were placed on the surgical platform where heat was controlled at 37 ° C and anesthetized using 3% isoflurane in oxygen through the nasal route. The hairs near the lower limb were cleaned, and the incision was made to expose the vein for scanning. Before the treatment, the mice were scanned using a flow probe (1 PRB; PR-series, Transonic system) of Power Doppler (Transonic, model T106) to record the blood flow in the femoral vein. Similarly, an Ultrasound imaging tool was assisted (Vevo LAZR-X; Fuji Film Visual Sonics, Inc.) for real time high resolution ultrasound imaging. After the first scanning, the animals were intravenously injected with saline, heparin, and KFeOx-NPs (10 mg/kg) treatment. Following an hour of treatment, subsequent imaging was performed following the generation of the thrombus model. The blood flow from the Doppler and the velocity were calculated and analyzed using Vevo LAB software. The veins were then dissected for histology. The blood was collected through cardiac puncture before sacrificing the mice, and the serum was separated. The standard protocol provided by the Mouse D2D (D-Dimer) ELISA kit manufacturer was followed to estimate the D-Dimer concentration (Catalogue No. EM0979).

### Application of NPs using a catheter

The medical-grade silicone catheter was used to check the capillary and implant-based action of KFeOx-NPs. The catheters with a width of ∼0.5 and ∼1.5 cm were dipped into the saturated solutions (overnight with constant stirring) of the respective treatment (EDTA and KFeOx-NPs) after the surface activation by plasma treatment. These were then air-dried to form a layer. The coated catheters were then incubated with blood with respective anticoagulants for 4 hours. These were then washed trice using PBS to remove the unbound blood cells. The catheter was then characterized using SEM before and after blood incubation. The proteins bound to the catheter were then determined using a protein estimation assay with a catheter length of ∼0.5 cm. Catheters were stained with Coomassie Brilliant Blue (CBB) for 40 minutes in shaking condition. Further washing with PBS, the OD of distaining was measured at 590 nm using a multiple reader (Synergy H1-Microplate reader) after 40 mins and plotted. Catheters with ∼1.5 cm were measured with the flow rate from one end to another to quantify the capillary action after and before blood incubation. 10% serum solution was used to pump from a catheter attached to a syringe using a syringe pump (SyringeONE) with a speed of 0.01 ml/min from a height of 19.5 cm. The time the catheter crossed 5 ml of serum solution was measured and plotted.

### Statistical Analysis

Statistical differences were calculated through one-way ANOVA analysis using OriginLab software. A value of p < 0.05 was considered statistically significant.

## Supporting information

https://docs.google.com/document/d/1O6YAh29MtaaevKIUsQ62UH0197v12816/edit?usp=drive_link&ouid=117484493709020649895&rtpof=true&sd=true

## Supporting Information

Supporting Information is available from the Wiley Online Library.

## Acknowledgments

S.M. is thankful to SERB (SRG/2023/000409), the Government of India, for providing financial support. The authors appreciate the staff of Dr. Lal Pathlab Varanasi, for human blood samples. DY is thankful to the Ministry of Human Resource Development (MHRD), Government of India, for her PMRF fellowship. LP is thankful to UGC; Pragya and Snehashis are thankful to CSIR, India for their research fellowships.

