## Supplementary material for "Potassium ferric oxalate nanoparticles prevent human blood clotting and thrombosis in a mouse model": https://docs.google.com/document/d/1O6YAh29MtaaevKIUsQ62UH0197v12816/edit?usp=drive_link&ouid=117484493709020649895&rtpof=true&sd=true

**Dynamic Light Scattering (DLS)**: A particle size analyzer employing the theory of dynamic scattering of light (S90, Malvern Zetasizer, U.K.) was utilized to measure the particle size. The solution of KFeOx-NPs was diluted ten times in Millipore water for the experiment. Similarly, the zeta potential was measured for the surface charge, which influences the stability.

**Ultraviolet Spectroscopy (UV):** The UV spectrophotometer was employed to measure the peak of KFeOx-NPs in the range of 200 -800 nm using UV spectrometer (BioSpectrometer, Eppendorf).

**High-Resolution X-Ray Diffraction (HR-XRD):** The XRD pattern of KFeOx-NPs was obtained using an X-ray diffractometer (Rigaku Miniflex 600, Jap Rigaku SmartLab 9kW Powder type (without $\chi$cradle), RIGAKU Corporation an). X-ray as the source of radiation, 40 kV tube voltage, and 15 mA current, with a step size of 0.05° at a 10−80° 2θ angle, were used to record the diffractograms.

**High-Resolution Scanning Electron Microscope (HR- SEM)**: The solid powder of KFeOx-NPs was imaged using SEM (Nova Nano SEM 450, FEI Company of USA (S.E.A.) PTE, LTD) at room temperature. Images were taken at magnifications of 25 Kx and voltage of 15 kV.

**High-Resolution Transmission Electron Microscope (HR- TEM):** 1mg/ml of KFeOx-NPs solution was used to coat the copper grid to investigate the shape, size, architecture, and surface morphology using TEM (Tecnai G2 20 TWIN, FEI Company of USA (S.E.A.) PTE, LTD).

**EDS:** The solid powder of KFeOx-NPs was imaged using SEM to detect the elements. Carbon, oxygen, iron, potassium, sulfur, calcium, and chlorine were detected using EDAX (Team Pegasus Integrated, EDS-EBSD with Octane Plus and Hikari Pro)

**Fourier Transform Infrared Spectroscopy (FTIR)**: The spectra of KFeOx-NPs and PVP were recorded in the range of 4000–500 $cm^{-1}$ by using a Fourier transform infrared spectrometer (Nicolet iS5, THERMO Electron Scientific Instruments LLC).

**Surface Area Measurement Facility (BET):** 200 mg solid powder of KFeOx-NPs was used to analyze the surface property by BET (BELLSORP MAX II & BELCAT-II, MicrotracBEL Corp.br> )

**Inductively Coupled Plasma Mass Spectrometry (ICP-MS):** The harvested organs were digested using nitric acid and diluted further to 5 ml for the analysis using ICPMS (Agilent 7800 ICP-MS mainframe, Agilent Technologies).

**Table S1**: BET of KFeOx-NPs

| Parameter | Values |
|---|---|
| **Average diameter of pores (nm)** | **2.5010** |
| **Mean pore diameter (nm)** | **0.5665** |
| **BET Surface area ($m^2/g$)** | **2.9297** |

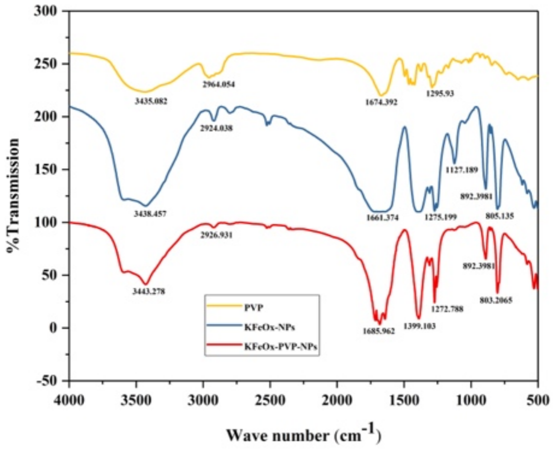


**Figure S1**: FTIR of PVP (yellow), KFeOx-NPs without PVP (blue), and KFeOx-NPs with PVP (red).

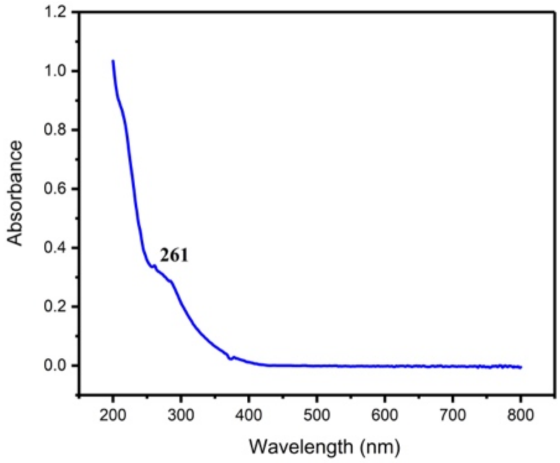


**Figure S2:** UV visible spectroscopy of KFeOx- NPs (without PVP)

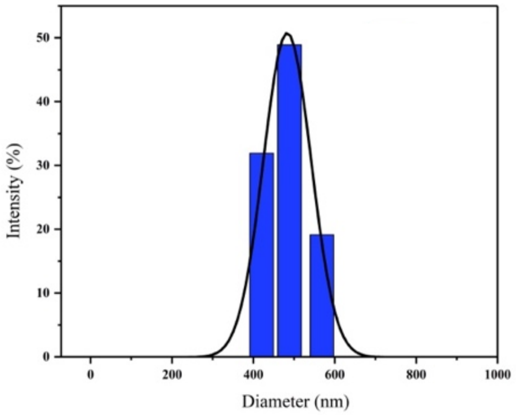


**Figure S3**: Dynamic light scattering (DLS) of KFeOx- NPs (without PVP)

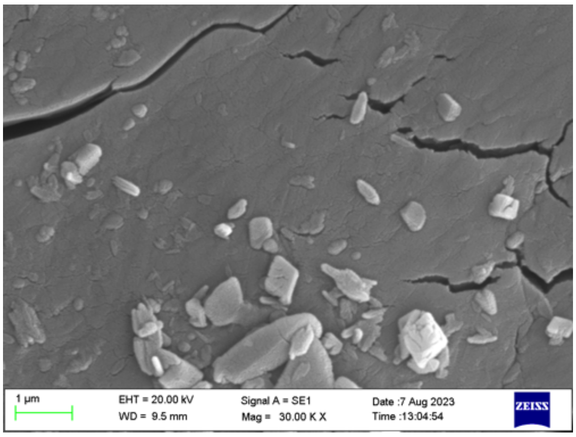


**Figure S4**: Scanning electron microscope (SEM) image of KFeOx- NPs (without PVP)

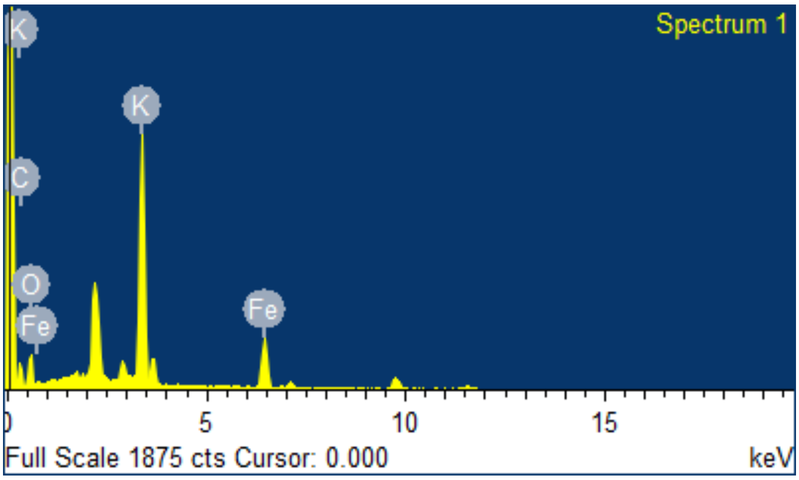


**Figure S5:** Energy-dispersive X-ray analysis (EDAX) of KFeOx- NPs (without PVP)

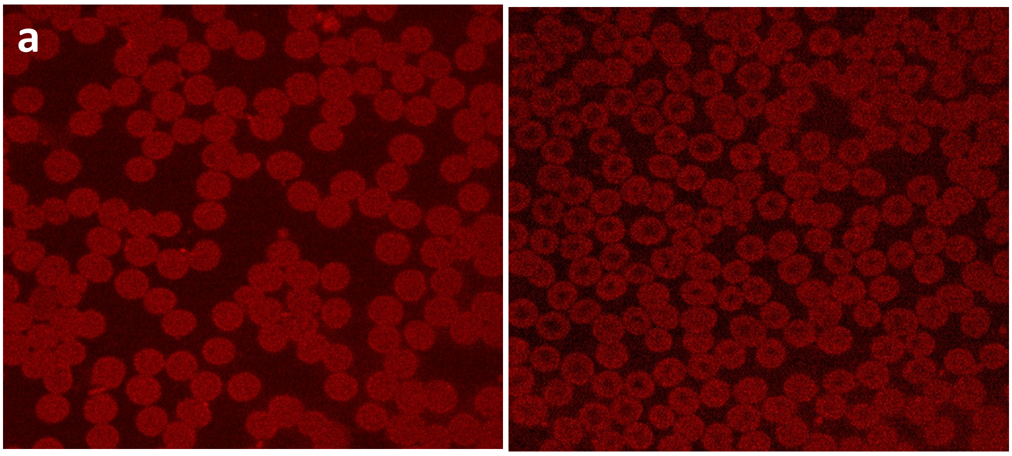


**Figure S6**: Confocal images of blood smear stained with rhodamine dye containing (a) EDTA, and (b) KFeOx-NPs

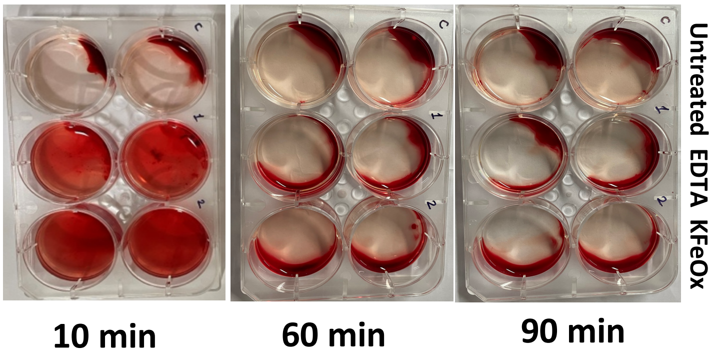


**Figure S7:** Dynamic clotting time assay with untreated and blood treated with EDTA and KFeOx-NPs

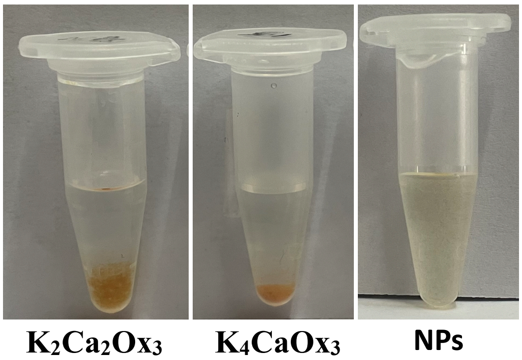


**Figure S8:** Pathological test confirming the presence of calcium oxalate formed after the interaction of calcium with the NPs

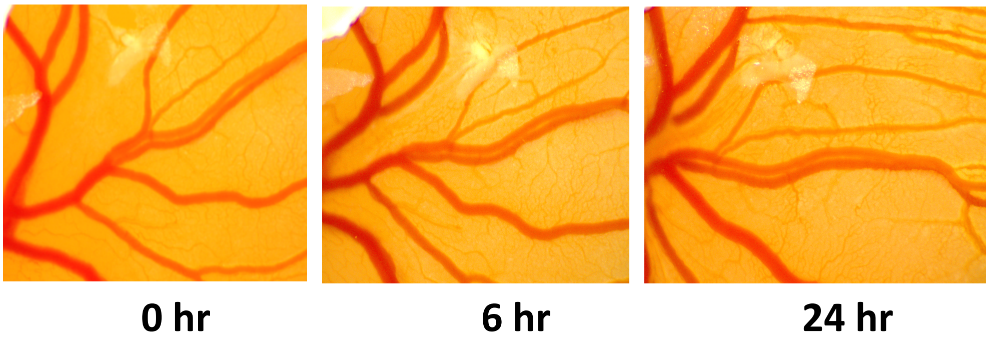


**Figure S9**: CAM images of KFeOx-NPs at 2 mg/ml till 24 hours of treatment

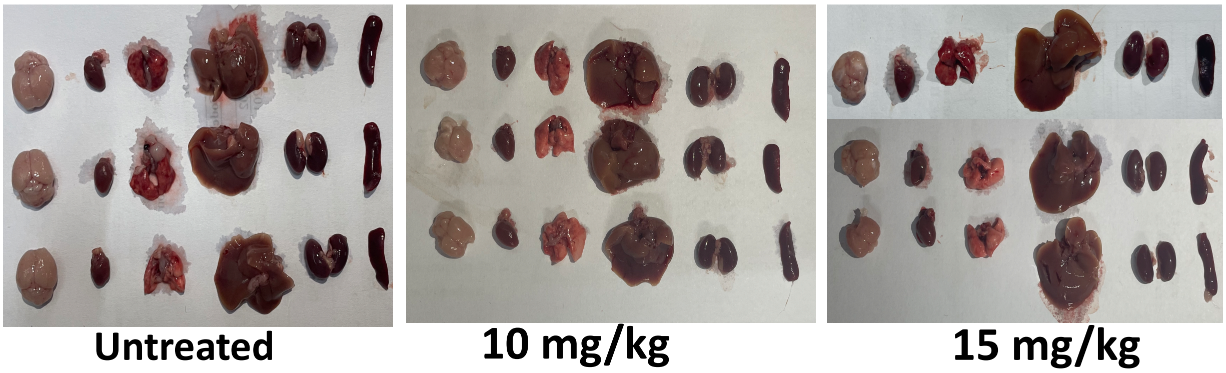


**Figure S10:** Optical images of organs (brain, heart, lungs, liver, kidney, and spleen) of animals of each group, namely untreated, 10mg/kg and 15 mg/kg of KFeOx-NPs treated

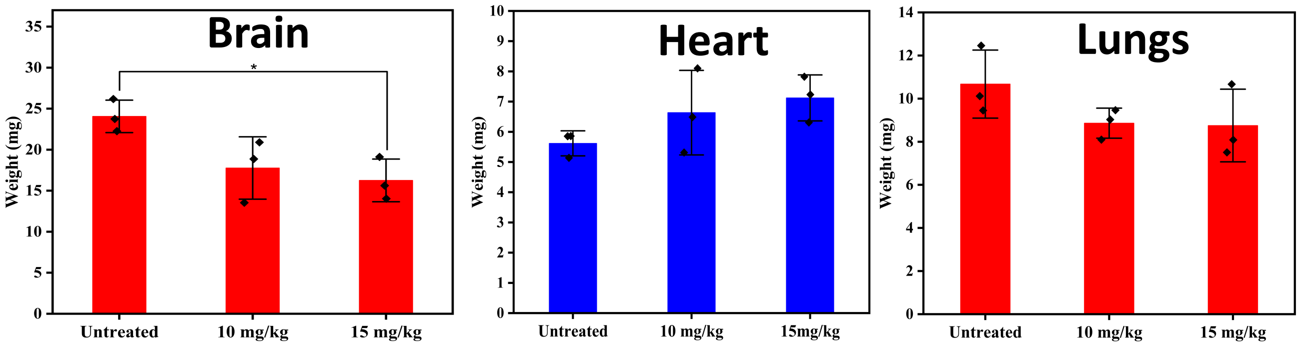


**Figure S11:** Weight of the organs (normalized with the individual body weight of mice) harvested after sacrificing the animals

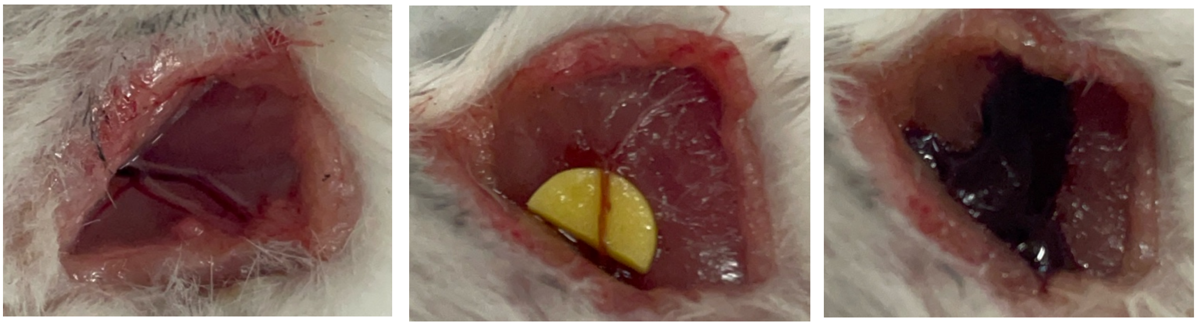

**Figure S12**: Ferric chloride-induced model for thrombosis. Showing femoral vein further induced with $FeCl_3$ dipped paper for 3-5 min, and then the incision of the vein

**Table S2:** Pathological analysis (CBC) for toxicity assessment

| Time | Sample | Neutrophil | Lymphocyte | Monocyte | Eosinophil | Basophil | IG |
|---|---|---|---|---|---|---|---|
| (Hrs) | | (%) | (%) | (%) | (%) | (%) | (%) |
| 0 | Untreated | 76.47 | 17.64 | 1.96 | 0.98 | 0 | 2.94 |
| | EDTA | 60.4 | 33.66 | 0.99 | 0.99 | 0 | 3.96 |
| | KFeOx-PVP-NPs | 61.38 | 33.66 | 0.99 | 0 | 0.99 | 2.97 |
| 6 | EDTA | 60.18 | 24.07 | 3.7 | 2.77 | 0.92 | 8.33 |
| | KFeOx-PVP-NPs | 50.5 | 29.7 | 10.89 | 3.96 | 1.98 | 2.97 |
| 24 | EDTA | 44.26 | 24.59 | 9.83 | 9.83 | 6.55 | 4.91 |
| | KFeOx-PVP-NPs | 58.87 | 17.75 | 6.54 | 14.95 | 1.86 | 0 |

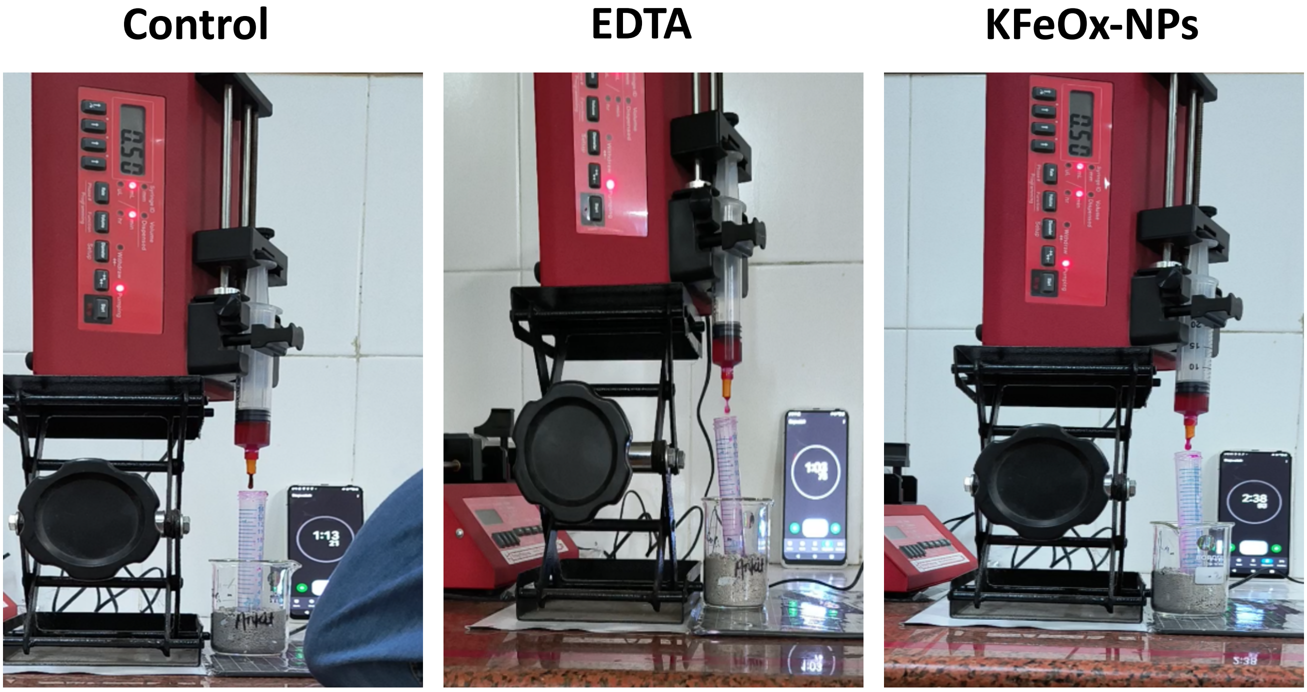

**Figure S13**: Flow property of the catheter incubated with the anticoagulants (control, EDTA, and KFeOx-NPs)

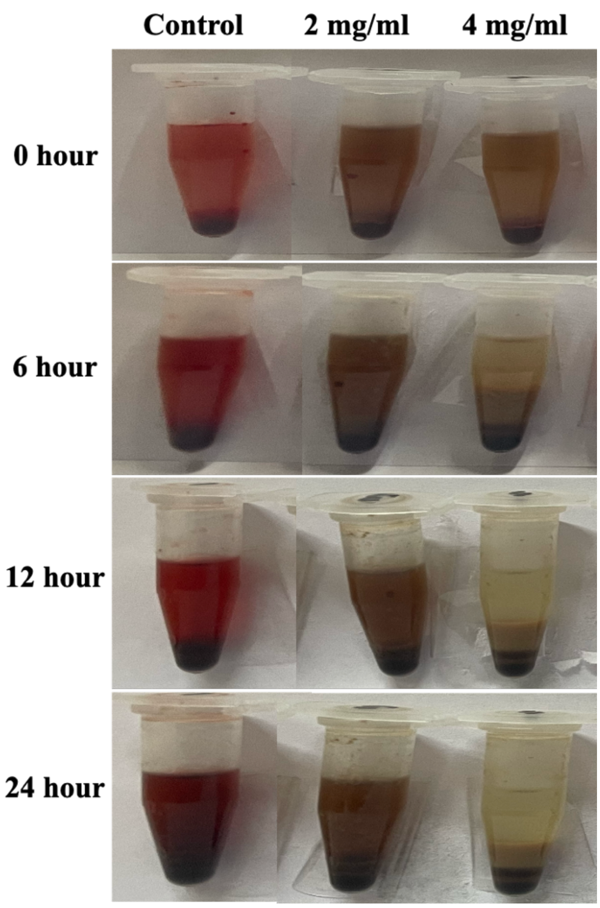


**Figure S14**: Thrombolytic assay of KFeOx-NPs with 2 and 4 mg/ml dose

**Video S1**: Prothrombin (PT) assay of blood containing EDTA and KFeOx-NPs
EDTA -
https://drive.google.com/file/d/1jf7lmPx1AmBqEhCeGUFjv0dIbezgHpSH/view?usp=share_link

KFeOx-NPs -
https://drive.google.com/file/d/1zEAPEAWZprTlQfR1waIIoi05_LiAHgF9/view?usp=share_link

**Video S2**: activated partial thromboplastin time (aPPT) assay of blood containing EDTA and KFeOx-NPs

EDTA - https://drive.google.com/file/d/1XavaCkAUp_5fSuZ0iA9LSD51klkJ8Lb8/view?usp=share_link

KFeOx-NPs - https://drive.google.com/file/d/1sMNXJOHLpJKmExgXmEA2U3MoIAopy7RV/view?usp=share_link

**Video S3**: Experimental video of tail bleeding assay for untreated, heparin and KFeOx-NPs treated groups.

Untreated - https://drive.google.com/file/d/1r8OfGLBmoIRjKUyG1-FzcUMSBdCpzikw/view?usp=share_link

Heparin - https://drive.google.com/file/d/1EO2b5nPxtJoXD0MzUjdJS9yxLa66JYs8/view?usp=share_link

KFeOx-NPs - https://drive.google.com/file/d/1o2zuvuWTKqrVJCwTE1lMIvMPPea3ZEBV/view?usp=share_link

**Video S4**: Blood flow captured from flow doppler before and after treatment using Vevo LAB software - https://docs.google.com/presentation/d/15wLRWa_6q6B4nxwyhgmqBEz-Mpjx6QnQ/edit?usp=share_link&ouid=117484493709020649895&rtpof=true&sd=true

**Video S5**: Kaolin clotting time assay of blood incubated with EDTA and KFeOx-NPs

EDTA - https://drive.google.com/file/d/1PofgjDGBlRXJxSBYuqsF8fHASZDcwmVB/view?usp=share_link

KFeOx-NPs - https://drive.google.com/file/d/1eiyTBJMDjtRuopS_CQm9LYMruuYuAej-/view?usp=share_link
